# Stable platelet production via the bypass pathway explains the long-term reconstitution capacity of hematopoietic stem cells

**DOI:** 10.1101/2024.06.23.599660

**Authors:** Shoya Iwanami, Toshiko Sato, Hiroshi Haeno, Longchen Xu, Keimyo Imamura, Jun Ooehara, Xun Lan, Hiromitsu Nakauchi, Shingo Iwami, Ryo Yamamoto

## Abstract

Precise understanding how hematopoietic stem cells (HSCs) differentiate in vivo is difficult because we can not trace the in vivo differentiation of HSCs. The single-cell transplantation assay and our own paired daughter cell assay of phenotypic HSCs has revealed the presence of HSCs with reconstitution capacity whose differentiation potential is restricted to the myeloid lineage (MySCs) and the presence of novel direct differentiation pathway form HSCs to MySCs (named myeloid bypass pathway). However, how HSCs differentiate in vivo during hematopoiesis has remained unclear since the paired daughter cell assay was performed partially ex vivo. Aiming to characterize HSCs, including the myeloid bypass pathway, we examined the kinetics of HSC differentiation using a mathematical model. We analyzed data from single-cell transplantation assays in which five blood cell lineages were followed successively after transplantation. An age-related skewing to the myeloid lineage was quantitatively indicated as the production of B cells reduced with age. Dependence on platelet bypass increased with aging and consistently high dependence was associated with long-term reconstitution capacity of the HSCs. Focusing on the ratio of chimerism among cell lineages, the characteristics of dependence on platelet bypass can be accurately determined by the ratio of erythrocyte to platelet chimerism at 8 weeks after transplantation. This new identification criterion is an indicator of long-term reconstitution of HSCs that does not rely on observation of long-term transplantation experiments. These findings reveal the novel characteristics of HSCs related to aging and stemness and highlight the importance of the bypass pathway in HSC differentiation.

**Significance statement:** In vivo differentiation of hematopoietic stem cells (HSCs) is important for understanding blood cell production. Here we investigated the differentiation kinetics of HSCs in single-cell transplantation assays by using a mathematical model. Our findings demonstrate the importance of the bypass pathway in platelet production for the long-term reconstitution capacity of HSCs. These results also suggest the utility of studying time changes in HSC differentiation. The new HSC characteristics and its detection criteria identified in this study are expected to be useful in understanding HSC diversity and aging.

## Introduction

Although it is known that mouse hematopoietic stem cells (HSCs) are found within the phenotypic CD150^+^CD34^-/low^Flt3^-^c-Kit^+^Sca-1^+^Lineage^-^ (CD150^+^CD34^-^KSL) cell population^1,2^, we are still unable to prospectively identify functional HSCs with 100% accuracy^3^. One powerful tool for characterizing the heterogeneity of HSCs is the single-cell transplantation assay. This tool evaluates the functional properties of transplanted cells by the duration of hematopoietic reconstitution and the blood cell lineages produced in the primary and secondary transplantations. In our previous studies using single-cell transplantation, we showed that HSCs producing all blood cell lineages can be distinguished with regard to the duration of reconstitution as long-term (LT-), intermediate-term (IT-), and short-term (ST-) HSCs^4,5^. Although single-cell transplantation experiments reveal the nature of the transplanted cells, this information is gained retrospectively and long-term observation is required to determine cell function.

We have previously shown using clonal transplantation assays that single HSCs demonstrate self-renewal and multipotency^6^. Furthermore, in the phenotypically defined HSC compartment, we found long-term repopulating cells with differentiation potential restricted to the myeloid (myeloid-restricted stem cells [MySCs]) at a single-cell level, which only generate megakaryocyte, erythrocyte, and megakaryocyte lineages, megakaryocyte and erythrocyte, or megakaryocyte (defined as common myeloid stem cells [CMSCs], megakaryocyte-erythroid stem cells [MESCs], and megakaryocyte stem cells [MkSCs], respectively)^4,7^, and identified a change in the frequency of MySCs with aging^5^. Paired-daughter cell assay combined with single-cell transplantation demonstrated the existence of a myeloid bypass pathway from HSCs; however, this assay was partially performed in vitro. Therefore, we cannot deny the possibility that this bypass pathway is an artificial phenomenon in vitro.

We hypothesized that mathematical modeling of the chimerism observed in peripheral blood (PB) could help to determine how hematopoiesis from HSCs depends on myeloid bypass using in vivo data alone. Using mathematical models to estimate cell differentiation kinetics allows us to understand the phenomena underlying time-related changes. For example, in a study analyzing the time course of leukemia cells in patients with chronic myeloid leukemia, investigators used the estimated decay rate of leukemic cells to identify target cells for therapeutic drugs and elucidate the mechanism of reactivation due to the emergence of resistant mutations after treatment^8^. In the multilineage differentiation of HSCs, mathematical models have been used to quantify the cell cycle changes associated with cell divisions and the subsequent bias in differentiation lineages^9^. Mathematical models have also been used to estimate differentiation pathways, which enabled comparison of the contribution of multiple pathways to blood cell production^10^. Theoretical analysis of the competitive proliferation of cells in transplantation assays and the reconstitution capacity of HSCs has also been conducted by use of a hierarchical cell differentiation model^11^.

Here, we sought to dissect LT-, IT- and ST-HSCs in the transplantation setting based on the reconstitution kinetics (chimerism) of the four lineages of blood cells (neutrophils/monocytes, erythrocytes, platelets, and B cells) using a mathematical model and to investigate how LT-, and IT-HSCs produce mature blood cell lineages via the myeloid bypass pathway. By applying mathematical modeling and clustering methodologies to functional chimerism datasets generated from 75 single-cell transplantation assays, we have proposed a new classification of four LT- and IT-HSC subsets. We found that HSCs biased toward platelets compared with erythrocytes at an early point after transplantation tended to be of the stable bypass subsets. Our large-scale datasets therefore provide a useful “atlas” of LT- and IT-HSC heterogeneity.

## Results

### Chimerism of donor cells as measured in HSC transplantation assays

In the transplantation assays used to evaluate the characteristics and capacity of cells involved in hematopoiesis, cells from labeled donor mice are transplanted into a recipient mouse. Then, the fraction of cells in PB derived from the transplanted cells, or *chimerism*, is measured as functional blood cell production (**Figure 1A**). In this study, we analyzed data from single-cell transplantation assays of phenotypic HSCs (pHSCs) performed in previous studies^4,5^ with the aim of elucidating the differentiation kinetics of HSCs. Chimerism values for neutrophils/monocytes, erythrocytes, platelets, and B cells from a total of 114 recipient mice were used (**Table 1**). Of the 114 mice, 81 were transplanted with young HSCs (8-12 weeks of age) and the remaining 33 were transplanted with aged HSCs (20-24 months of age). The class of transplanted single cells was defined according to the duration of hematopoietic reconstitution with chimerism values measured according to a defined threshold during primary and secondary transplantation. A total of 29, 35, and 39 single cells were classified as long-term HSCs (LT-HSCs), intermediate-term HSCs (IT-HSCs), and short-term (ST-HSCs), respectively. The remaining 11 single cells were not evaluated by secondary transplantation and could not be distinguished as LT-HSC or IT-HSC (i.e., LT/IT-HSCs). Here, we used the chimerism data from the first transplantation only, so that LT-HSCs, LT/IT-HSCs, and IT-HSCs were considered as a single population named LT-HSCs in our mathematical model for the following analysis. Complete blood cell count (CBC) was measured in 30 of the transplantations for the experiments that fit the criteria for data use (**Materials and Methods**). Focusing on the chimerism dataset, there was no clear age-related difference in the time course of chimerism for any of the HSC classes (**Figure 1B**). Comparing LT-HSC, LT/IT-HSC, and IT-HSC chimerism at week 24 between young and aged mice, there was a significant trend toward a decrease with age in B cells only (p-value was 0.0134, **Figure 1C**). As for the definition of HSC classes, there were clear differences in chimerism between LT- or IT-HSC and ST-HSC, but the differences between LT-, IT/LT-, and LT-HSC were unclear and the chimerisms were highly heterogeneous (**Figure 1B**). See **Table 1, Materials and Methods**, and the original studies^4,5^ for details on the experiment and the dataset.

**Table 1.** Summary of dataset from transplantation assay.

| Transplanted<br>KuO<sup>+</sup> cells | Single cells (N = 114 recipient mice) |  |  |  | Bone marrow<br>cells (N = 10<br>mice) |
| --- | --- | --- | --- | --- | --- |
| Data type | Class |  |  |  | Recipient |
|  | LT-HSC in modeling<br>(LT-HSC:LT/IT-HSC:IT-HSC) |  | ST-HSC |  |  |
|  | Young | Old | Young | Old |  |
| Chimerism | 53<br>(23:5:25) | 22<br>(6:6:10) | 28 | 11 | 10 |
| CBC | 10<br>(3:1:6) | 7<br>(3:1:3) | 6 | 7 | — |

**Figure 1.**
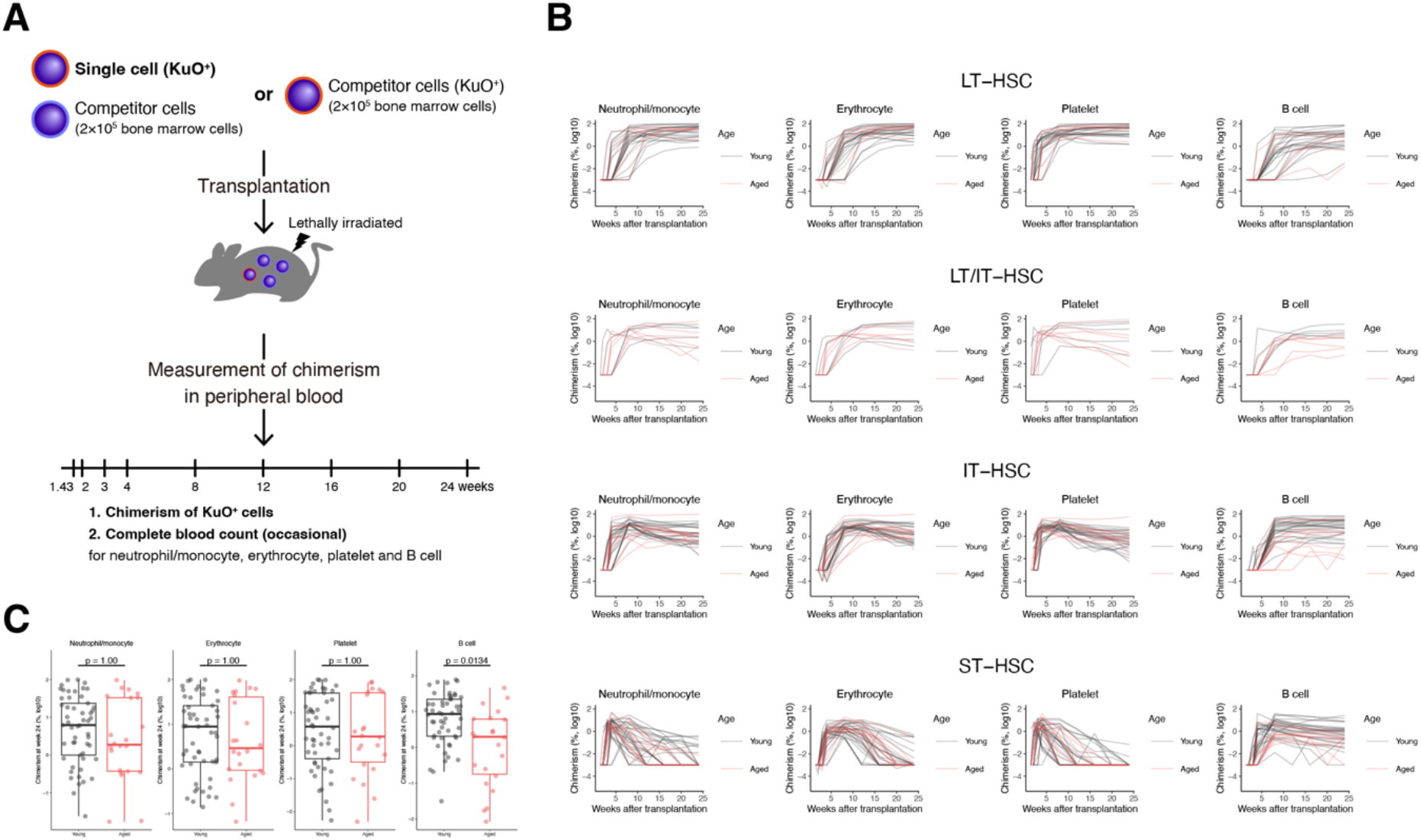
Single-cell transplantation assay and scheme of mathematical model of HSC differentiation. **(A)** Schematic representation of the single-cell transplantation assay of pHSCs. Single pHSCs obtained from donor mice expressing Kusabira-Orange (KuO) were transplanted into lethally irradiated recipient mice along with 2 × 10^5^ bone marrow cells. In all transplantation assays, the percentage of cells derived from donor mouse cells in each blood cell lineage in the PB (chimerism) was measured over 24 weeks. In part of the transplantation assays, complete blood cell count (CBC) was measured at the same time. Details of the transplantation assays and data collection were described in previous studies^4,5^. **(B)** The time-course change in chimerism obtained in single-cell transplantation assays for each class, LT-HSCs, LT/IT-HSCs, IT-HSCs, and ST-HSCs, defined by reconstitution capacity. The black and red lines indicate chimerism of single cells obtained from young and aged mice, respectively. Data below the detection limit are displayed with chimerism as 0.001%. **(C)** Comparison of chimerism from LT-HSCs, LT/IT-HSCs, and IT-HSCs for each blood cell lineage at 24 weeks after transplantation in the single-cell transplantation assay with aging. P-values were calculated by the Student’s t-test for the mean of young and aged HSCs with Bonferroni correction.

### A mathematical model of the direct myeloid lineage production pathway from HSCs explains blood cell production in vivo

The fraction of cells in the PB observed in a single-cell transplantation assay reflects not only the capacity of a single donor cell but also competition with simultaneously transplanted competitor cells and the recipient mouse’s own cells. Chimerism must be evaluated throughout the time course taking into account the context in which the data are collected. We used a mathematical model describing the hierarchical differentiation of HSCs using ordinary differential equations (**Eqs. (1-7)**) to capture the time-course changes in data from the single-cell transplantation assays (**Figure 2A and B**). Data fitting and numerical simulation with this model were performed to quantitatively understand the capacity of transplanted single cells and the factors contributing to changes in chimerism. Our model describes blood cell production by hierarchical differentiation of self-renewing LT-HSCs (i.e., LT-HSCs, LT/IT-HSCs, and IT-HSCs) and ST-HSCs (**Figure 2A**). In addition, we assumed that there was a differentiation pathway from LT-HSCs to myeloid progenitors that was not via ST-HSCs. This pathway is defined as the *myeloid bypass pathway*, as revealed in previous studies by MySCs, whose differentiation potential is restricted to myeloid cells^4,7^. The chimerism of each lineage in the transplantation assay is assumed to be based on the number of mature cells produced from progenitor cells (**Eqs. (8-9)**). The amount of each cell population produced at each time point from the upper population is defined as *influx*, which is an indicator of the differentiation potential of HSCs (**Figure 2B**). In particular, the degree of dependence on the myeloid bypass pathway can be assessed by comparing the influx to myeloid progenitor cells from the two pathways described in the mathematical model. See the **Supplementary Text** for more details on the mathematical model.

**Figure 2.**
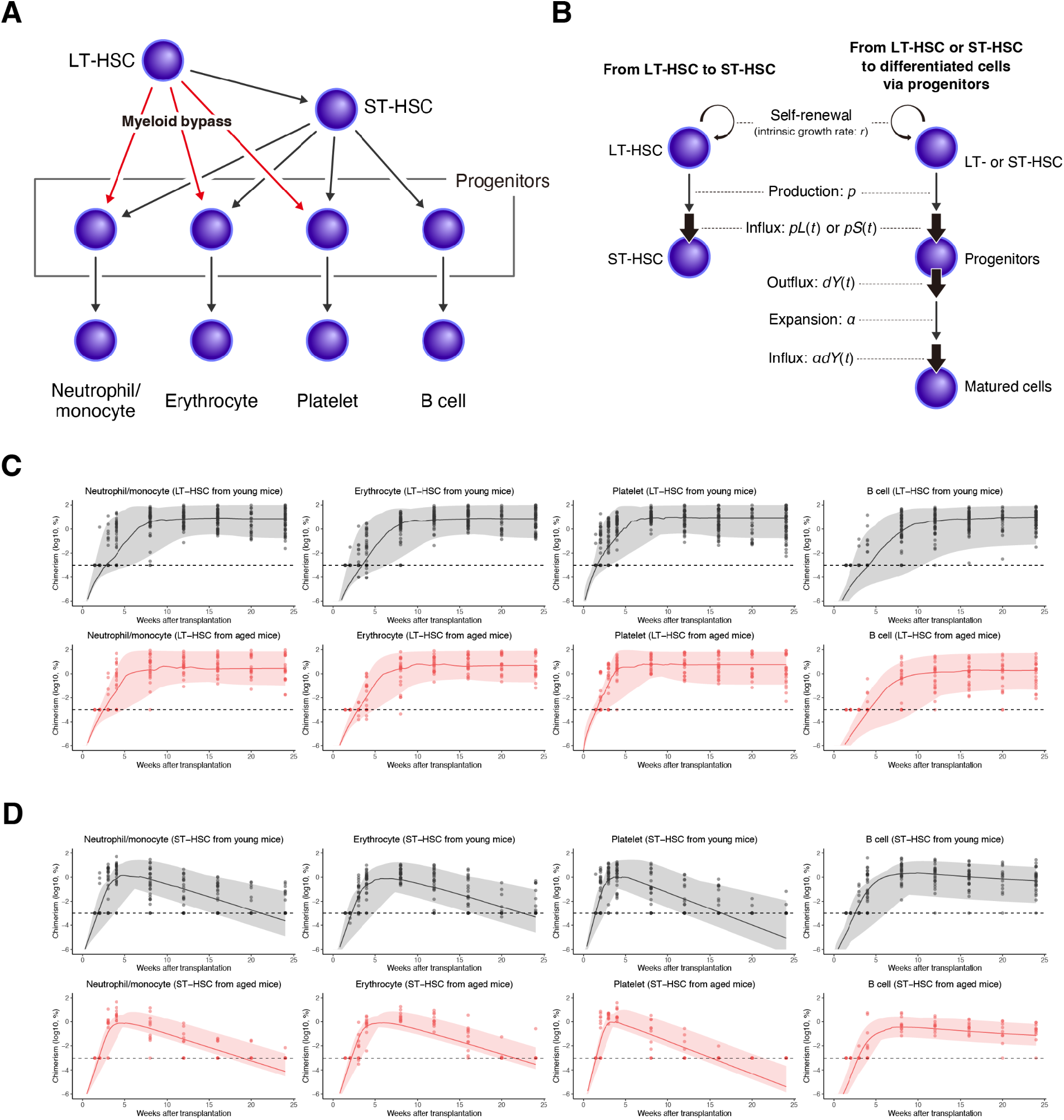
Summary of the data fitting to chimerism data using a mathematical model. **(A)** Overview of the hierarchical structure of the differentiation model. The pathway for differentiation from LT-HSCs to progenitor cells that does not go through ST-HSCs is termed the *myeloid bypass pathway* and is indicated by the red arrows. **(B)** Proliferation by self-renewal and blood cell production by differentiation of each cell population from LT-HSCs to ST-HSCs (left) and from LT-HSCs or ST-HSCs to mature cells via progenitor cells (right) as described in the mathematical model. The number of cells flowing into the lower cell population as a result of cell differentiation from the cell population in the upper layers is defined as *influx*. **(C-D)** Value of chimerism (dots) and predictions calculated by the mathematical model in transplantation assays of single cells obtained from young **(C)** and aged **(D)** mice. The solid lines show the median of the mathematical model’s predictions and the filled area represents the minimum and maximum of the chimerism data calculated with estimated individual parameters. See **Supplementary Text** for the calculation of prediction intervals.

To assess the capacity of transplanted single HSCs, we estimated the parameters of the mathematical model by using the chimerism and CBC of neutrophils/monocytes, erythrocytes, platelets, and B cells in the single-cell transplantation assay (**Figures 2, S1 and S2**). The percentage of cells on the recipient side with transplantation of bone marrow cells labeled with Kusabira-Orange (KuO) (**Figure S3**) were obtained independently for this study and used for data analysis. All data were simultaneously used to estimate the parameters of the mathematical model with a nonlinear mixed-effects model. The values of the model parameters estimated as population parameters and individual parameters are listed in **Tables S1** and **S2**, respectively. The chimerism calculated using the estimated parameters captures the overall trend in the data from the single-cell transplantation assay.

### Age-related changes in HSC capacity cause a myeloid shift due to a decrease in B cell production

The values of model parameters for individual single cells estimated from the data fitting allowed us to assess age-related changes in HSC capacity. In our mathematical model, the self-renewal capacity of HSCs is corresponds to the intrinsic growth rate of LT-HSCs and ST-HSCs, *r*_*L*_ and *r*_*S*_, respectively. Comparison of the values of these parameters indicated that the self-renewal capacity of LT-HSCs remained the same or slightly decreased with age (the self-renewal capacity of aged LT-HSCs was 0.738 times that of young LT-HSCs, **Table S1** and **Figure 3A**), whereas that of ST-HSCs did not change (the self-renewal capacity of aged ST-HSCs was 1.11 times that of young ST-HSCs, **Table S1** and **Figure 3B**). Focusing on the differentiation of HSCs into progenitor cells, the capacity to produce ST-HSCs from LT-HSCs, *p*_*LS*_, increased with age (**Figure 3A**), but the capacity to produce progenitor cells of each blood cell lineage from ST-HSCs, *p*_*SN*_, *p*_*SN*_, *p*_*SE*_, and *p*_*SP*_, decreased (**Figure 3B**). In particular, the capacity to produce the myeloid lineage (neutrophils/monocytes, erythrocytes, and platelets) was greatly reduced (*p*_*SN*_, *p*_*SE*_, and *p*_*SP*_ were 0.322, 0.597, and 0.379 times lower in aged HSCs, **Table S1** and **Figure 3B**). Myeloid bypass tended to increase platelet production capacity, *p*_*LP*_, which was 2.75 times higher in aged HSCs (**Table S1** and **Figure 3A**). On the other hand, neutrophil/monocyte and erythrocyte production, *p*_*LN*_ and *p*_*LE*_, tended to be low even in young LT-HSCs and decreased further with age (**Figure 3A**). The values of production capacity from HSCs to progenitors, *p*_*LN*_, *p*_*LE*_, *p*_*LP*_, *p*_*SB*_, *p*_*SP*_, *p*_*SE*_, and *p*_*SN*_, represent the ratio of weekly production to the size of the progenitor population. The bypass pathways to produce neutrophils/monocytes and erythrocytes from LT-HSCs, *p*_*LE*_, *p*_*LP*_, were suggested to be largely inactive because their values were less than 10^−6^ per week even for young HSCs. The estimated HSC capacity and their age-related changes are summarized in **Figure 3C**.

**Figure 3.**
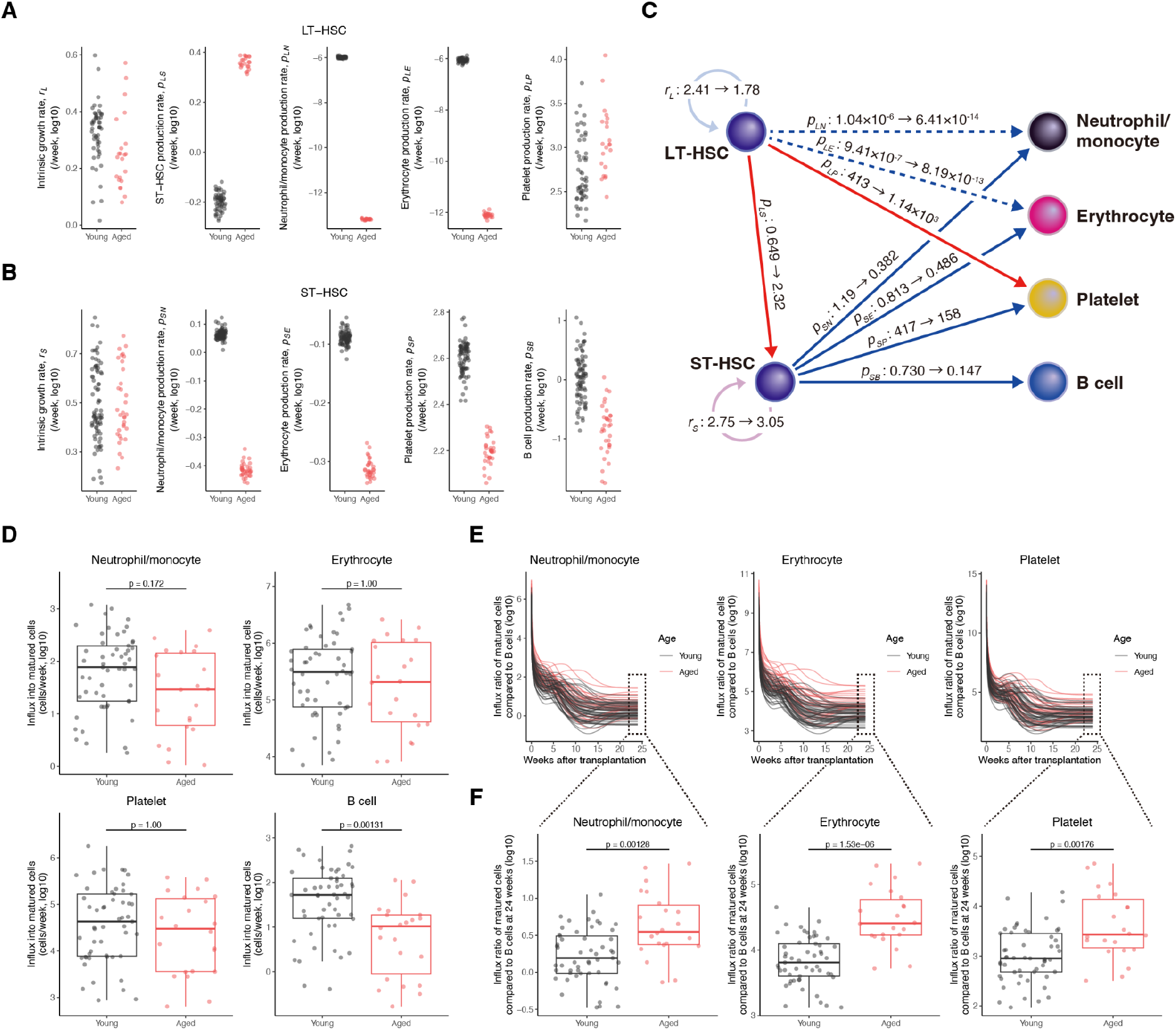
Age-related changes in estimated HSC capacity. **(A)** Age-related changes in the estimated intrinsic growth rate, *r*_*L*_, ST-HSC production rate, *p*_*LS*_, platelet production rate, *p*_*LP*_, erythrocyte production rate, *p*_*LE*_, and neutrophil/monocyte production rate, *p*_*LN*_, of LT-HSCs. **(B)** Age-related changes in the estimated intrinsic growth rate, *r*_*S*_, B cell production rate, *p*_*SB*_, platelet production rate, *p*_*SP*_, erythrocyte production rate, *p*_*SE*_, and neutrophil/monocyte production rate, *p*_*SN*_, of LT-HSCs. **(C)** Graphical summary of age-related changes in HSC capacity corresponding to **(A** and **B)**. Red and blue arrows indicate increases and decreases, respectively. The intensity of the color of the arrow indicates the magnitude of the changes. Few progenitor cells are produced in the pathway depicted by the dashed line. **(D)** Age-related changes in estimated production (i.e., influx) from progenitors into mature cells in each cell lineage at 24 weeks after transplantation (i.e., *α*_*i*_*d*_*i*_*Y*_*i,D,k*_(24) (*i* = *N, E, P* and *k* = *n*_*L*1_, …, *n*_*L*75_) and *d*_*B*_*E*_*B*2,*D,k*_ (24) (*k* = *n*_*L*1_, …, *n*_*L*75_)). **(E-F)** Production of each lineage of myeloid cells relative to the production of B cells at each time after transplantation (i.e., *α*_*i*_*d*_*i*_*Y*_*i,D,k*_ (*t*)/ *d*_*B*_*E*_*B*2,*D,k*_ (*t*) (*i* = *N, E, P* and *k* = *n*_*L*1_, …, *n*_*L*75_), **(E)**) and at 24 weeks after transplantation (i.e., *α*_*i*_*d*_*i*_*Y*_*i,D,k*_ (24)/ *d*_*B*_*E*_*B*2,*D,k*_ (24) (*i* = *N, E, P* and *k* = *n*_*L*1_, …, *n*_*L*75_), **(F)**). P-values in **(D)** and **(F)** were calculated by the Student’s t-test for the mean of young and aged HSCs with Bonferroni correction.

Although changes in the self-renewal capacity of HSCs with aging and the degree of differentiation as a transition between populations have been quantified, we need to examine changes in the number of cells produced as a result of differentiation to assess the differentiation potential of HSCs. To quantify how age-related changes in LT-HSCs affect blood cell production, we compared the amount of each cell lineage produced from donor cells at each time point calculated with estimated individual parameters. Comparing the influx of mature cells from young and aged HSCs, we found that the production of B cells, *d*_*B*_*E*_*B*2,*D*_(24), significantly decreased with age (the p-value was 0.00131), whereas the production of myeloid cells, *α*_*i*_*d*_*i*_*Y*_*i,D*_ (*i* = *N, E, P*), remained the same (**Figure 3D**). The ratio of production of the three myeloid cell lineages to that of B cells as representative of lymphocytes, *α*_*i*_*d*_*i*_*Y*_*i,D*_(*t*)/*d*_*B*_*E*_*B*2,*D*_(*t*) for *i* = *N, E, P*, tended to increase with age in all lineages throughout transplantation (**Figure 3E**). These ratios at week 24, *α*_*i*_*d*_*i*_*Y*_*i,D*_(*t*)/*d*_*B*_*E*_*B*2,*D*_(24) for *i* = *N, E, P*, were significantly increased in all lineages (**Figure 3F**), which is consistent with the myeloid skewing of HSCs with aging reported in another study^12^. The above results suggest that, at least under transplantation conditions, a myeloid shift in HSCs with aging was confirmed as the absolute number of mature blood cells. This skewing toward myeloid differentiation was caused by decreased lymphoid production rather than by increased myeloid production.

The HSC transplantation assay has been used to evaluate the self-renewal and differentiation potential of HSCs. The magnitude of chimerism is sometimes regarded as the strength of self-renewal or differentiation of HSCs; however, given the mechanism of the transplantation system, it may also reflect the differentiation potential of the blood cell lineage being observed and the characteristics of competing cells. The estimated self-renewal rate of LT-HSCs, which is expressed as an intrinsic growth rate in our model, *r*_*L*_, was slightly positively correlated with chimerism at 24 weeks in B cells only (**Figure S4A**). However, there was much variability and it seems unlikely that a linear relationship explains this relationship (**Figures S4B and S4C**). The number of LT-HSCs in the competitor cells, which encompasses not only the number relative to transplanted single cells but also relative capacity, showed a slight negative correlation except for B cells (**Figure S4C**). However, there was large variation in this metric as well (the second column from right in **Figure S4A**). The strongest correlation was found between the intensity of B cell production from ST-HSC, *p*_*SB*_, and B cell chimerism (**Figure S4C**), which may still explain at most 50.7% of B cell chimerism (**Figure S4B**). To understand the capabilities of HSCs, it is necessary to evaluate the kinetics of multiple strains of chimerism rather than chimerism of a single lineage at a single time point.

### Changes in the degree of dependence of platelets on myeloid bypass represent differences in the time course of mature cell production

The dependence on the bypass pathway in differentiation from LT-HSCs into myeloid lineages was defined as the fraction of influx into progenitor cells that was not via ST-HSCs in the overall influx in single-LT-HSC transplantations (i.e., *p*_*Li,D*_*L*_*D*_(*t*)/(*p*_*Li,D*_*L*_*D*_ (*t*) + *p*_*Si,D*_*S*_*D*_(*t*)), **Figure 4A**). Since the calculated blood cell production from donor cells was found to be roughly steady-state at 24 weeks after transplantation (**Figure S5**), we compared the fraction at week 24 as a property of HSCs in post-reconstitution of hematopoiesis. The degree of dependence on myeloid bypass at week 24 was higher for platelets than for neutrophils/monocytes and erythrocytes (**Figures 4A and B**). The platelet dependency on the bypass pathway at week 24 was significantly increased in aged HSCs compared with young HSCs (the p-value was 0.0142), whereas that for neutrophils/monocytes and erythrocytes was suggested to decrease with age (the p-values were 4.03 × 10^−48^ and 4.69 × 10^−^, respectively), and were quite low, less than 10^−5^, even in young HSCs (**Figure 4A** and **B**). In the process of blood cell reproduction from a single cell, the degree of dependence on platelet bypass changed with time after transplantation (**Figure 4A**). Particularly notable was the decrease in the degree of dependence on platelet bypass up to 10 weeks after transplantation in some HSCs regardless of the degree of dependence at week 24 or the age of the transplanted HSCs (**Figures 4A and C**). Based on the difference in dependency on platelet bypass, HSCs can be divided into two categories: HSCs with a degree of dependence on the bypass pathway of less than 20% at any time point were defined as “dropped”, and all other HSCs were defined as “stable”. To also take into account the dependency at week 24 compared above, we defined HSCs with dependency lower than 50% at 24 weeks after transplantation as “low”, and the rest as “high” (**Figure 4C**). The majority of LT-HSCs were stable-high (22 of 29 LT-HSCs), suggesting a correlation between long-term reconstitution capacity and dependence on platelet bypass (**Figure 4D** and **Table 2**). Surprisingly, aged HSCs defined as “dropped” tended to recover the extent of bypass dependency (9 of 10 aged HSCs defined as dropped), whereas young HSCs defined as dropped did not recover the extent (5 of 16 young HSCs defined as dropped). HSCs defined as dropped were IT-HSCs (11 of 16 HSCs defined as dropped including LT/IT-HSCs). Aged HSCs defined as dropped-high were not LT-HSCs (1 of 9 aged HSCs defined as dropped-high), suggesting that the increase in dependence on platelet bypass seems to occur in relation to age, not stemness (**Figure 4D**).

**Table 2.** Platelet bypass dependency in single-cell transplantation.

| Age | Young (N = 53) |  |  | Aged (N = 22) |  |  |
| --- | --- | --- | --- | --- | --- | --- |
| Class | LT-HSC | LT/IT-HSC | IT-HSC | LT-HSC | LT/IT-HSC | IT-HSC |
| Stable-High | 17 | 3 | 4 | 5 | 2 | 5 |
| Stable-Low | 3 | 1 | 9 | 0 | 0 | 0 |
| Dropped-High | 2 | 1 | 2 | 1 | 4 | 4 |
| Dropped-Low | 1 | 0 | 10 | 0 | 0 | 1 |

**Figure 4.**
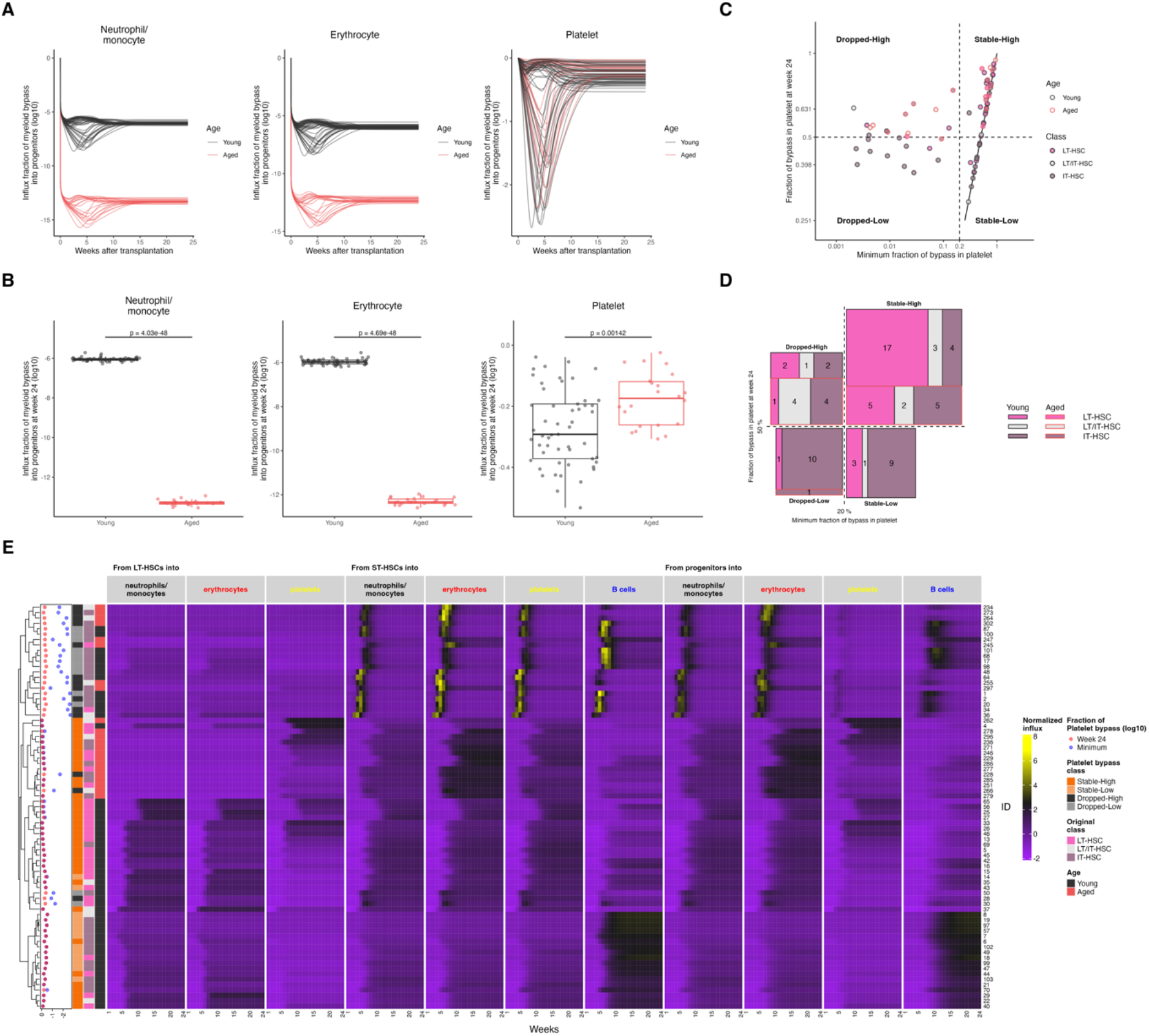
Dependence on the bypass pathway in myeloid cell production and HSC skewing in differentiation. **(A-B)** Dependency on the bypass pathway in each lineage of myeloid cells at each time after transplantation (i.e., *p*_*Li,D,k*_*L*_*D,k*_(*t*)/(*p*_*Li,D,k*_*L*_*D,k*_ (*t*) + *p*_*Si,D,k*_*S*_*D,k*_ (*t*)) (*i* = *N, E, P* and *k* = *n*_*L*1_, …, *n*_*L*75_), **(A)**) and at 24 weeks after transplantation (i.e., *p*_*Li,D,k*_*L*_*D,k*_ (24)/(*p*_*Li,D,k*_*L*_*D,k*_ (24) + *p*_*Si,D,k*_*S*_*D,k*_ (24)) (*i* = *N, E, P* and *k* = *n*_*L*1_, …, *n*_*L*75_), **(B)**). **(C)** Correlation of minimum dependency of platelet production on the bypass pathway in each transplantation assay and dependency at 24 weeks after transplantation. Solid vertical and horizontal lines show the dependency at 50% and 20%, respectively. **(D)** Relationship between classes of HSCs defined by platelet bypass and aging, and classes defined by reconstitution capacity. The number of HSCs of the type indicated by the color corresponds to the size of the rectangle, which corresponds to **(C)** and which are summarized in **Table 2. (E)** Hierarchical clustering of production of each cell population. The values at 24 weeks after transplantation (red) and the minimum (blue) for the fraction of platelet bypass in the platelet progenitor production are shown next to the dendrogram. The left side of the production heatmap shows the class defined by platelet bypass, the class defined by the duration of reconstitution after transplantation, and of the transplanted HSCs. P-values in **(B)** were calculated by the Student’s t-test for the mean of young and aged HSCs with Bonferroni correction.

To investigate how the different time-dependent characteristics of the dependency on platelet bypass affected the hematopoietic potential of HSCs, we calculated continuous time-course changes of influx from upper to lower cell populations with estimated model parameters (**Figure S6**) and compared the weekly production of the four lineages of individual HSCs after single-cell transplantation (**Figure 4E**). In our mathematical model, there were influxes from HSCs into progenitor cells and from progenitor cells into mature cells. First, we normalized by the maximum value for all mice for each influx to compare all influxes equally. Next, to consider skewing in blood cell production within individuals, we normalized the influx in each individual so that it followed a normal distribution with a mean of 0 and a variance of 1. Hierarchical clustering using the distance of normalized influx clearly divided HSCs into two major groups defined as dropped and stable. The dropped group showed high mature cell production of neutrophils/monocytes, erythrocytes, and B cells from week 3 to 15, whereas the stable group had relatively high production after week 8 (4 large columns on the right in **Figure 4E**). In addition, the stable-low group produced more B cells than the other lineages (bottom of first column on the right for the individuals with light orange for the platelet bypass class in **Figure 4E**). These comparisons suggested that differences in the differentiation pathway to platelets could explain the overall skewing in the productive capacity of HSCs (i.e., platelet bypass dependency affects the production of mature blood cell lineages).

### Platelet bypass is correlated with platelet-biased potential compared to erythrocyte

To date, pHSCs have been classified by combining reconstitution duration data in primary and secondary transplants (LT, IT, or ST) and the extent of differentiation skewing to neutrophils/monocytes, B cells, and T cells at week 20-24 in the primary transplants (myeloid-biased, lymphoid-biased, or balanced). We wondered whether the dependence on platelet bypass that we found to be associated with reconstitution capacity and aging could be defined from the value of the data in the transplantation assay. We first aimed to characterize these large PB chimerism datasets by use of a clustering approach.

Hierarchical clustering of PB chimerism data from 75 single-cell transplantations revealed three distinct clusters, Cluster I, II, and III (**Figure 5A and Materials** and **Methods**). The three clusters were divided into two main groups according to the time of myeloid predominance. In Cluster I, the chimerism of myeloid cells was predominant in the latter 12 to 24 weeks after transplantation. In contrast, Clusters II and III had a predominance of myeloid cells in the first half of the transplantation, from 4 to 12 weeks. Clusters II and III were distinguished by the predominance of lymphocytes at 12-24 weeks. This clustering could capture a particular lineage skewing in chimerism at a certain time point after transplantation. Interestingly, our analysis separated LT- and IT-HSCs into two major populations, Cluster I and Cluster II/III, respectively. Cluster I mainly consisted of HSCs defined as stable-high for platelet bypass, whereas Clusters II and III consisted of HSCs defined as low and high for platelet bypass. This suggests that the class defined by the degree of dependency on platelet bypass reflects not only the reconstitution capacity of HSCs, but also the skewing of chimerism as time change of relative values between blood cell lineages.

**Figure 5.**
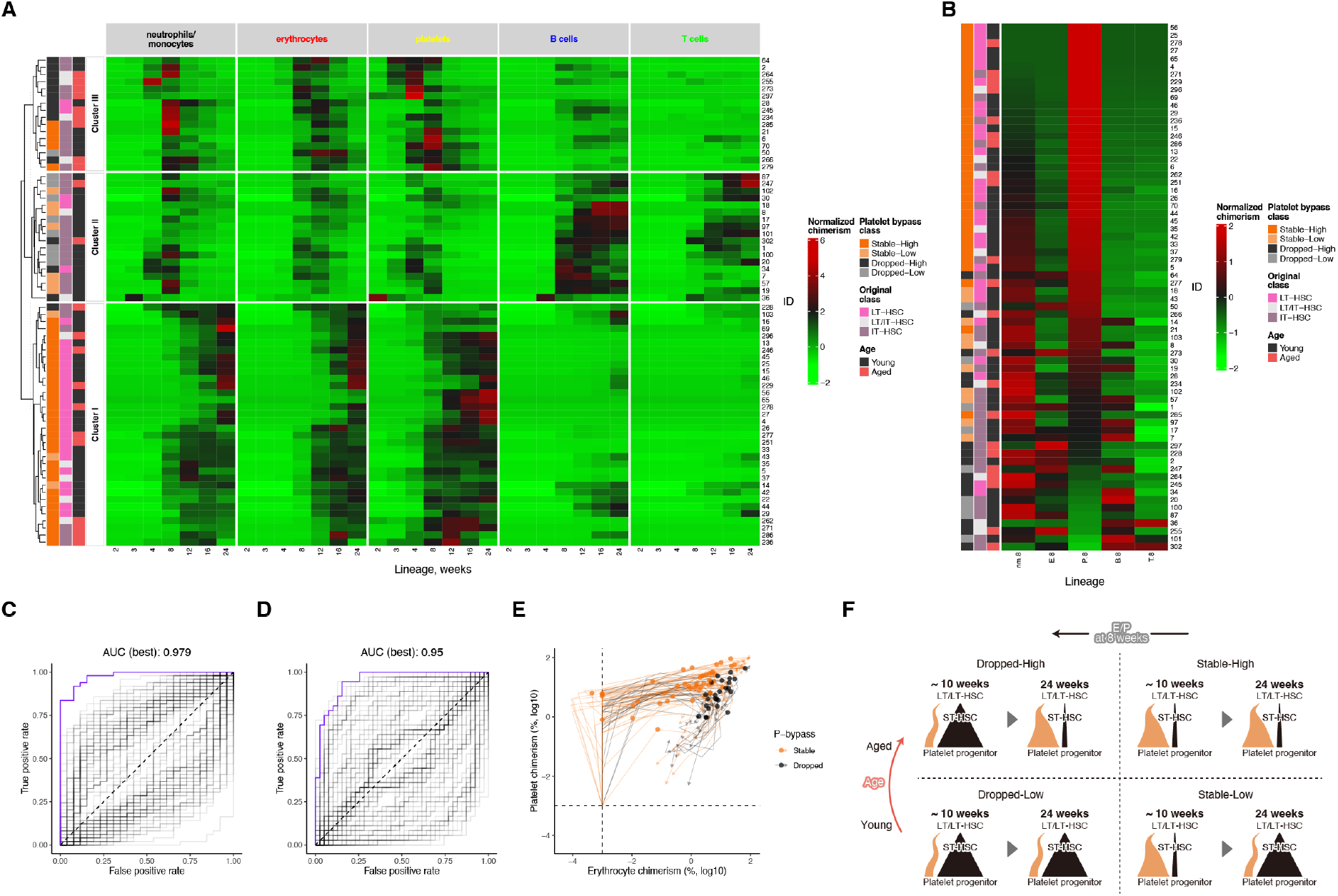
HSCs in the stable class of platelet bypass have platelet-biased potential. **(A)** Hierarchical clustering of chimerism of each cell lineage observed in the transplantation assay. Each transplanted individual was divided into three groups by the k-means method. **(B)** Normalized chimerism at 8 weeks after transplantation with HSCs classified as LT-HSC or IT-HSC. The individuals are sorted in decreasing order using the normalized chimerism value of the platelet. The class defined by platelet bypass, the class defined by the duration of reconstitution after transplantation, and age for transplanted HSCs are shown next to the heatmap in **(A)** and **(B). (C-D)** ROC curves obtained for classification of two classes, stable and dropped **(C)** and stable-high and others **(D)**, defined by platelet bypass dependency with ratio of chimerism as an explanatory variable. Each line was obtained by changing the threshold for the combination of lineages of chimerism data and observation time used for classification, and group assignment. The purple line shows the ROC curve obtained by using the ratio of erythrocyte to platelet chimerism at 8 weeks after transplantation, when the area under the curve (AUC) was highest. **(E)** Time change of erythrocyte and platelet chimerism values and class of platelet bypass. Erythrocyte and platelet chimerism in transplantation assays classified into stable and dropped classes based on platelet bypass dependency is shown by orange and black lines. The direction of the arrow indicates the passage of time after transplantation. The dots show chimerism values for platelets and erythrocytes at 8 weeks after transplantation. **(F)** Illustration of the transition in platelet production and the distinction between each class defined by platelet bypass. The orange and black bands indicate production from the bypass pathway and the pathway via ST-HSCs, respectively.

We investigated the relationship between the platelet bypass and chimerism data at each time point after transplantation based on platelet dominance using relative values of chimerism at 2, 3, 4, 8, 12, 16, and 24 weeks. To assess skewing in chimerism for the five observed lineages, we normalized the chimerism values to follow a distribution with a mean of 0 and variance of 1 within each individual. The values were then sorted in descending order by normalized chimerism of platelet values indicating platelet predominance and examined in relation to other blood cell lineages and classes defined by platelet bypass (**Figures 5B and S7**). This analysis revealed that relative values of platelet chimerism at 4 to 24 weeks correlated highly with the stable-high class for platelet bypass (**Figures 5B and S7**), especially suggesting that the values at 8 weeks separate the stable and dropped classes regardless of whether the class was high or low (**Figures 5B**). In addition, the stable and dropped classes were observed to correlate inversely with relative values of chimerism in lineages other than platelets at 8 weeks after transplantation (**Figure 5B**). These findings demonstrate the possibility of classifying HSCs with classes of platelet bypass that encompass the reconstitution capacity and the bias in the blood cell production of HSCs by examining skewing in chimerism at a given time point in transplantation assays.

The analysis of the pHSC population so far suggests that platelet bypass is important for functional HSCs. Therefore, we searched for features that could classify the stable and dropped classes of platelet bypass that could roughly distinguish functional LT-HSCs from IT-HSCs. In particular, we comprehensively investigated whether the ratio of chimerism could separate the stable from the dropped class, since the skewing in HSC blood cell production and chimerism data was suggested to be relevant to the classes. For all combinations of the five cell lineages and observation times, we evaluated the accuracy and specificity for the stable and dropped classifications based on the chimerism ratio of two lineages with varying thresholds. Based on the area under the curve (AUC) of the receiver operating characteristic (ROC) curve, we found that the ratio of platelet to erythrocyte chimerism at 8 weeks after transplantation best separated the stable from the dropped class (AUC was 0.979, shown in purple line in **Figure 5C**). The classification of stable-high, to which most LT-HSCs belonged, and other classes was equally possible (AUC was 0.95, shown in purple line in **Figure. 5D**). The time variation of the combination of erythrocyte and platelet chimerism values differed between the stable and dropped classes with a clear difference at week 8 (circles in **Figure 5E**), whereas platelet values were relatively higher in the stable class and erythrocyte values were higher in the dropped class. Taken together, these data indicate that platelet bypass is strongly correlated with skewing toward platelet compared with erythrocyte chimerism. The data also imply that the dependency on platelet bypass defined by mathematical modeling can be estimated by the chimerisms ratio of platelet versus erythrocyte at 8 weeks before long-term observation of chimerism (**Figure 5F**).

## Discussion

Here, we have elucidated how and when transplanted HSCs use the platelet bypass pathway through mathematical analysis of PB chimerism datasets. We performed this analysis without using the paired-daughter cell assay and without analyzing cell division in the bone marrow (BM). Almost all LT-HSCs showed stable dependency on platelet bypass throughout the analysis (up to 24 weeks), and we observed increased dependency on platelet bypass with aging. This finding is consistent with previous reports showing an increased number of platelet-biased HSCs, or MkSCs, with aging^5,13^. On the other hand, we found that the dependency on neutrophil/monocyte bypass and erythrocyte bypass decreased with aging and that the absolute amounts of production via the bypass pathway were low. When we compared the classes of dependency on platelet bypass, we could identify HSCs with stable functioning of platelet bypass (AUC of ROC curve was 0.979) with use of a simple formula calculating the ratio of erythrocyte to platelet chimerism at 8 weeks. Stable platelet production via the bypass pathway is a requirement for long-term blood cell renewal as LT-HSCs (especially young LT-HSCs), and this new property can be determined by use of transplantation assays of relatively short duration. By investigating cell populations in the BM after HSC transplantation, we can clarify the differentiation kinetics of HSCs with greater certainty.

The percentage of cells derived from donor cells in PB in transplantation has been used by itself to assess reconstitution capacity and differentiation skewing, but we found that those values did not always reflect the estimated HSC self-renewal and differentiation potential. Given that our model also includes the effects of competitive proliferation with competitor cells, the mathematical modeling analysis gives a more precise assessment of the differentiation potential of HSCs. In particular, we showed a relative predominance of myeloid production by aged HSCs, with a decrease in B cell production. Several transplantation assays, transcriptome analysis, and tracking by fluorescent protein and genetic barcoding have shown a bias of the HSCs themselves toward myeloid differentiation and myeloid-dominant blood cell production in mice and humans^9,10,12–18^. Our estimated and calculated results are also consistent with the results of these previous studies. It is a new finding that, at least in the transplantation setting, the absolute amount of myeloid production does not change.

Previously, we elucidated a differentiation pathway of HSCs using paired-daughter cell assay, where two daughter cells after one division were transferred into separate mice to compare in vivo differentiation potential^4^. This is the only way to compare the differentiation potential of the two daughter cells in vivo; however, a single cell needs to divide in vitro. Our new approach, combining in vivo data with mathematical modeling, has successfully quantified how this pathway could work in a transplantation setting. Our mathematical model, which captures the kinetics of blood cell production, predicted time-dependent changes in dependence on the bypass pathway and revealed changes in HSC reconstitution capacity with age.

So far the mechanisms underlying myeloid (especially platelet) bypass remain to be elucidated. Direct comparison between daughter cells after one division of HSCs may help to identify these mechanisms. However, we cannot estimate whether the HSC divides via myeloid bypass or not. In other words, we cannot analyze which HSCs will pass through myeloid bypass. One approach would be to analyze the gene expression of many daughter cells when HSCs divide, but their gene expression cannot be mapped to their reconstitution capacity. Exhaustive screening is also difficult, but the model derived in this study should make this possible. Our simple calculation can identify platelet bypass at 8 weeks after transplantation. If we focus on the HSCs that differentiate to give rise to platelets via platelet bypass, we can detect candidate molecules to promote or inhibit platelet bypass. Using this method, we will be able to obtain candidate genes that will affect platelet bypass dependency after CRISPR screening.

In conclusion, the approach developed here aids in the quantitative understanding of HSC differentiation and redefines how HSC competence is assessed. The dependence on platelet bypass and its time- and age-related changes highlighted by our findings may contribute to a better understanding of the stemness and differentiation of HSCs.

### Limitations

The differentiation hierarchy of the model from LT-HSCs to progenitors is debatable. It is not clear whether ST-HSCs always differentiate from LT-HSCs. In this model, the production of ST-HSCs from LT-HSCs in competitor cells results in single ST-HSCs that are eliminated, representing their short blood cell production. Blood cell production via multipotent progenitors, common myeloid progenitors, and common lymphoid progenitors can all be interpreted as rounding to production from ST-HSCs to progenitors. The results for platelet bypass were obtained by focusing only on the bypass pathway from LT-HSCs. The existence of bypass pathways from more differentiated stages, such as ST-HSCs in this case, in the sense that they do not go through a common progenitor cell, is not ruled out and these pathways may play an important role.

In transplantation assays, the percentage of donor cells in the PB cannot be measured if the cells are not engrafted. Given that we observed blood cell production from a very small number of HSCs, i.e., single cells; it is possible that there is a stochastic failure to engraft. Therefore, the differentiation kinetics of HSCs estimated in this study may be the result of a survival bias of those that happen to be reconstitutive in single-cell transplantation. Even in light of these limitations, our results captured the dynamics of hematopoiesis during HSC reconstitution and provided new insights, especially for platelet bypass.

## Materials and methods

### Dataset of chimerism and CBC in single-cell transplantation

The longitudinal chimerism data from single-cell transplantation were previously published^4,5^. Chimerism was measured as the fraction of KuO^+^ cells in each cell lineage—neutrophil/monocyte, erythrocyte, platelet, B cell, and T cell—after transplantation of young and aged KuO^+^ phenotypically defined HSCs (CD150^+^CD41^-^CD34^-^KSL, CD150^+^CD41^+^CD34^-^KSL or CD150^-^CD34^-^KSL, pHSCs) into lethally irradiated mice. A total of 2 × 10^5^ BM cells were transplanted along with single HSCs in these assays. For the estimation of our model parameters, we used the data from single-cell transplantation in which transplanted single cells were classified as long-term (LT-) HSCs, intermediate-term (IT-) HSCs, and short-term (ST-) HSCs (**Table 1, Figures 1B and S1**). The data with measurements less than three points in at least one blood cell lineage were excluded. The chimerism data of T cells were excluded for parameter estimation to avoid having to estimate a large number of parameters. CBCs measured in a subset of experiments in which the chimerism data met the above criteria were also used to estimate parameters for the mathematical model (**Table 1** and **Figure S2**).

### Recipient chimerism in BM cell transplantation

To consider the effect of the percentage of the recipient’s original cells on erythrocyte and platelet chimerism in single-cell transplantation assays, transplantations of KuO-labeled BM cells into lethally irradiated mice were conducted as additional experiments (**Table 1**). As in the single-cell transplant experiment, the fraction of donor and recipient cells in erythrocytes and platelets in the PB after transplantation was measured. The percentage of recipient-derived cells was used for fitting the mathematical model (**Fig. S3**).

### Mathematical model for chimerism and CBC in single-cell transplantation and chimerism in BM cell transplantation

In the single-cell transplantation assays conducted in previous studies^4,5^, a single phenotypically defined HSC (donor cell) was sorted from donor mice which continuously express the fluorescent protein, KuO, and transplanted into recipient mice with BM cells (competitor cells) to rescue the initial blood cell production. We used the following mathematical model of the differentiation dynamics from HSCs into neutrophils/monocytes (*i* = *N*), erythrocytes (*i* = *E*), platelets (*i* = *P*), and B cells (*i* = *B*) during the single-cell transplantation for donor cells (*j* = *D*), competitor cells (*j* = *C*), and recipient cells (*j* = *R*):

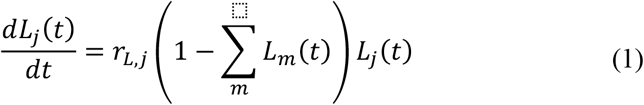

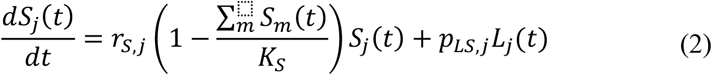

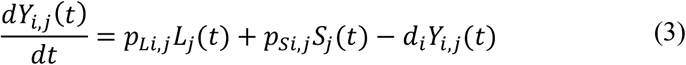

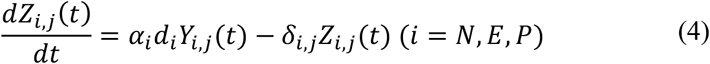

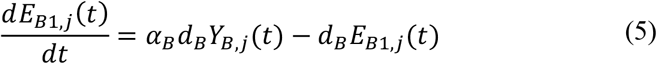

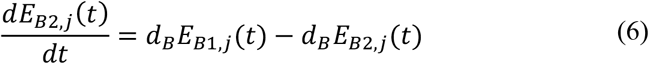

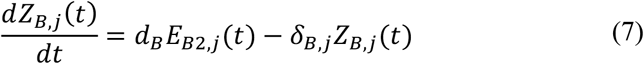

Here time *t* represents the number of weeks after transplantation, and the time when the one-cell transplant was performed is *t* = 0. In this model, LT-HSC, *L*_*i*_(*t*), and ST-HSC, *S*_*i*_(*t*), and progenitor cell, *Y*_*i,j*_(*t*), populations are calculated as cell densities. LT-HSCs and ST-HSCs proliferate by self-renewal at intrinsic growth rates *r*_*L*_ and *r*_*S*_, respectively. *K*_*S*_ is the carrying capacity of ST-HSCs. ST-HSCs differentiate from LT-HSCs at rate *p*_*LS,j*_. Progenitor cells of lineage *i* differentiate from LT-HSCs and ST-HSCs at rate *p*_*Li,j*_ and *p*_*Si,j*_, respectively. For B cells, *p*_*LB,j*_ = 0 because the bypass pathway is not considered. Myeloid progenitors migrate out of the population by differentiation at rate *d*_*i*_, producing *α*_*i*_*d*_*i*_*Y*_*i,j*_ (*t*) mature cells per week. In B cells, we hypothesized that an eclipse phase, *E*_*B*1,*j*_ and *E*_*B*2,*j*_, exists in the differentiation from progenitor cells into mature cells to represent the delayed onset of chimerism in the single-cell transplantation. The number of cells of each lineage measured in PB at time *t* is represented by *Z*_*i,j*_ (*t*), and mature cells die at rate *δ*_*i,j*_.

In the single-cell transplantation assay, cells derived from a single cell express the fluorescent protein KuO, which distinguishes them from competitor cells and recipient cells. Since recipient mice express only Ly5.2, whereas competitor cells express both Ly5.1 and Ly5.2, we can distinguish them in the neutrophil/monocyte and B lineages. Therefore, chimerism measured for each lineage *i* at time *t* in a single-cell transplantation assay, *F*.(*t*) can be expressed as follows using **Eqs. (1-7)**:

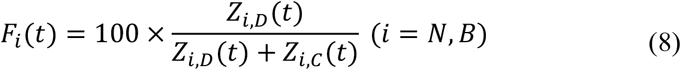

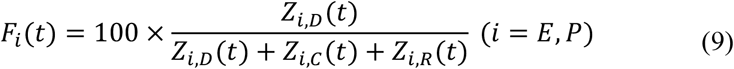

The CBC for each lineage *i* at time *t, T*_*i*_ (*t*), is described as follows:

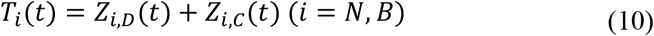

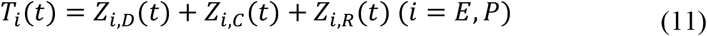

For more details about the mathematical modeling, see **Supplementary Text**.

### Parameter estimation by nonlinear mixed-effect model

Using a nonlinear mixed-effects model, the parameters of the mathematical model were estimated from data obtained in single-cell and BM cell transplantation assays. Data fitting was implemented using MonolixSuite2023R1^19^. For more details about the parameter estimation, please see **Supplementary Text**.

### Clustering of influx and PB chimerism datasets

Hierarchical clustering was performed and figures are plotted in R (http://www.r-project.org) using the ComplexHeatmap package^20,21^. The influx calculated by the mathematical model (**Eqs. (1-7**)) with estimated parameters was normalized by dividing by the maximum of each influx value to range from 0 to 1. It was then normalized to have a mean of 0 and variance of 1 within individual mice. Chimerism values in PB were normalized from the original values to have a mean of 0 and variance of 1 within individuals. Hierarchical clustering was performed using complete linkage.

## List of Supplemental Materials

- Table S2 (TableS2.csv)
- Figures S1-S8 (FigureS1.pdf - FigureS8.pdf)

## Acknowledgements

The authors thank Jennifer Holmes for providing language editing support.

## Author contributions

**Conceptualization**, S. Iwami and R.Y.; **Methodology**, S. Iwanami, T.S., H.H., L.C., X.L. and S. Iwami; **Software**, S. Iwanami, T.S. and L.C.; **Validation**, S. Iwanami, H.H. and S. Iwami; **Formal Analysis**, S. Iwanami and T.S.; **Resources**, H.N. and R.Y.; **Data Curation**, J.O. and R.Y.; **Investigation**, S. Iwanami, T.S., H.H., S. Iwami and R.Y.; **Writing – Original Draft**, S. Iwanami, K.I., S. Iwami and R.Y.; **Writing – Review & Editing**, S. Iwami and R.Y.; **Visualization**, S. Iwanami and T.S.; **Supervision**, S. Iwami and R.Y.; **Project Administration**, S. Iwami and R.Y; **Funding Acquisition**, S. Iwanami, H.H., S. Iwami and R.Y.

## Data availability

The original data of the transplantation assays will be shared by the lead contact upon request after publication.

## Supplemental information

### Supplementary figures

**Figure S1.**
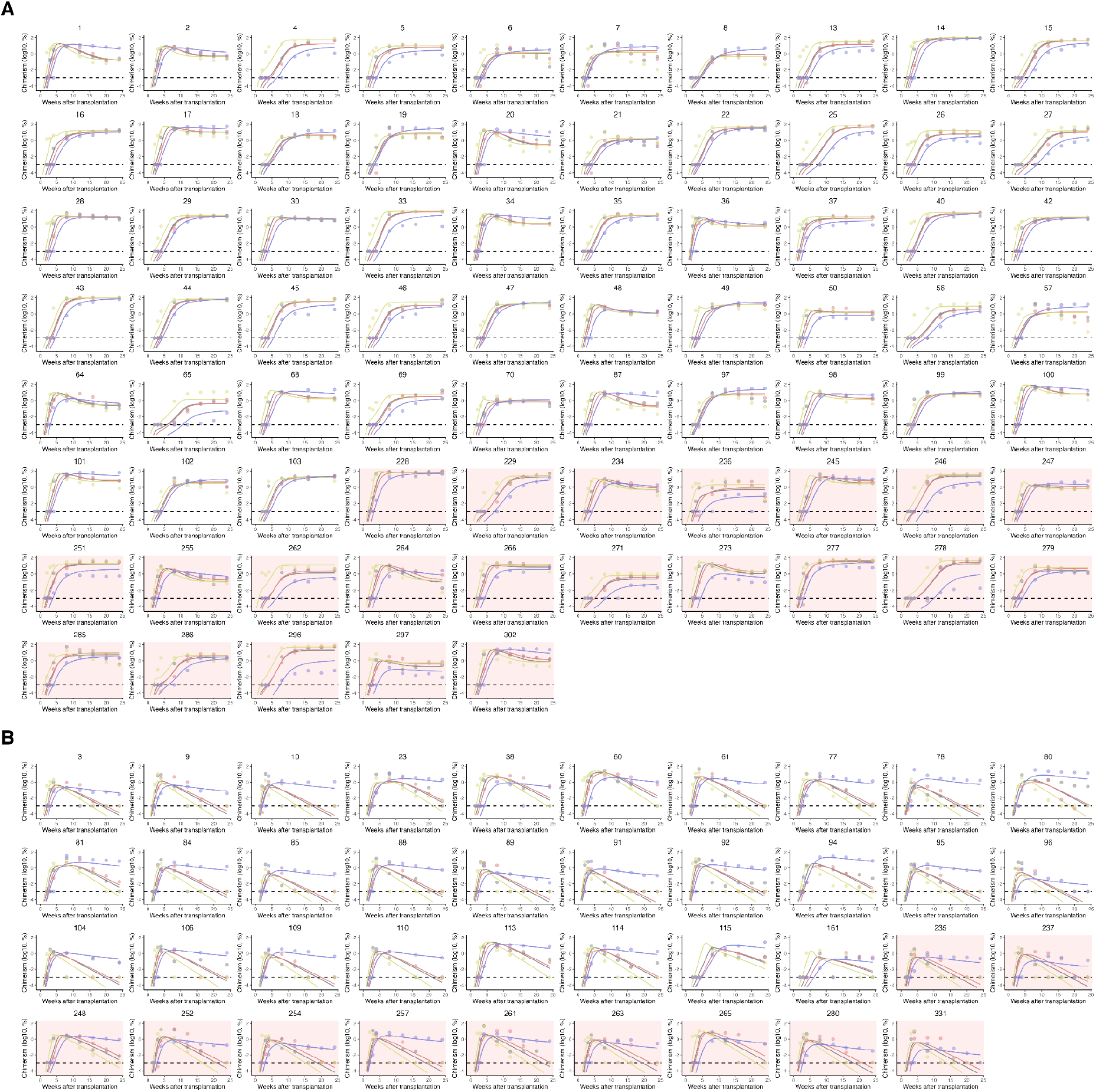
Individual chimerism data in the single-cell transplantation assay and best-fit model. **(A-B)** Individual chimerism data from the single-cell transplantation assay (dots) and expected values of the mathematical model calculated with best-fit parameters (lines). Dots and lines in gray, red, yellow, and blue correspond to neutrophils/monocytes, erythrocytes, platelets, and B cells, respectively. Panels with white and pink backgrounds are experiments in which young and aged HSCs were transplanted, respectively. **(A)** and **(B)** are experiments in which transplanted HSCs were classified as LT-HSC and ST-HSC, respectively.

**Figure S2.**
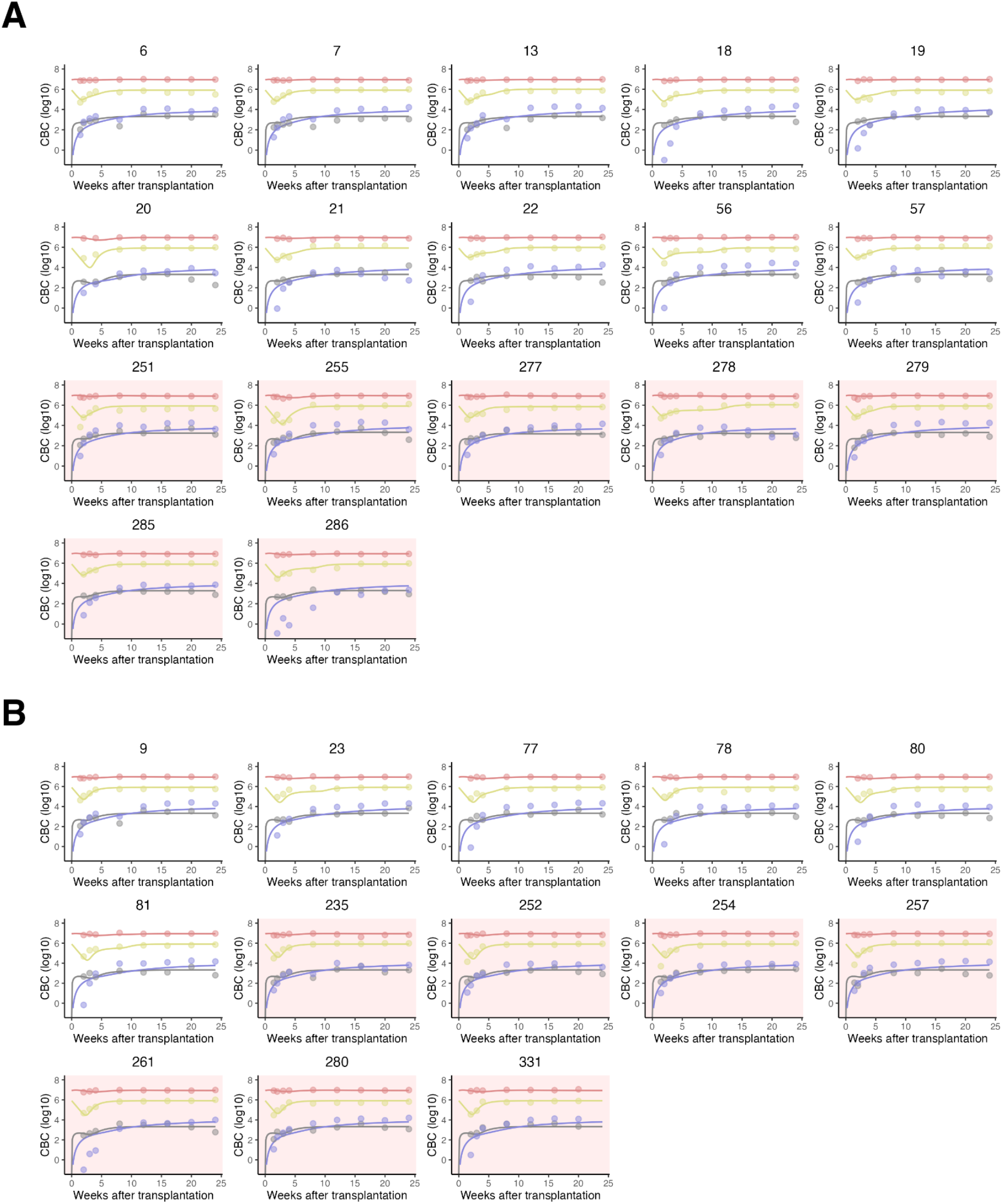
Individual CBC data in the single-cell transplantation assay and best-fit model. **(A-B)** Individual CBC data from the single-cell transplantation assay (dots) and expected values of the mathematical model calculated with best-fit parameters (lines). Dots and lines in gray, red, yellow, and blue correspond to neutrophils/monocytes, erythrocytes, platelets, and B cells, respectively. Panels with white and pink backgrounds are experiments in which young and aged HSCs were transplanted, respectively. **(A)** and **(B)** are experiments in which transplanted HSCs were classified as LT-HSC and ST-HSC, respectively.

**Figure S3.**
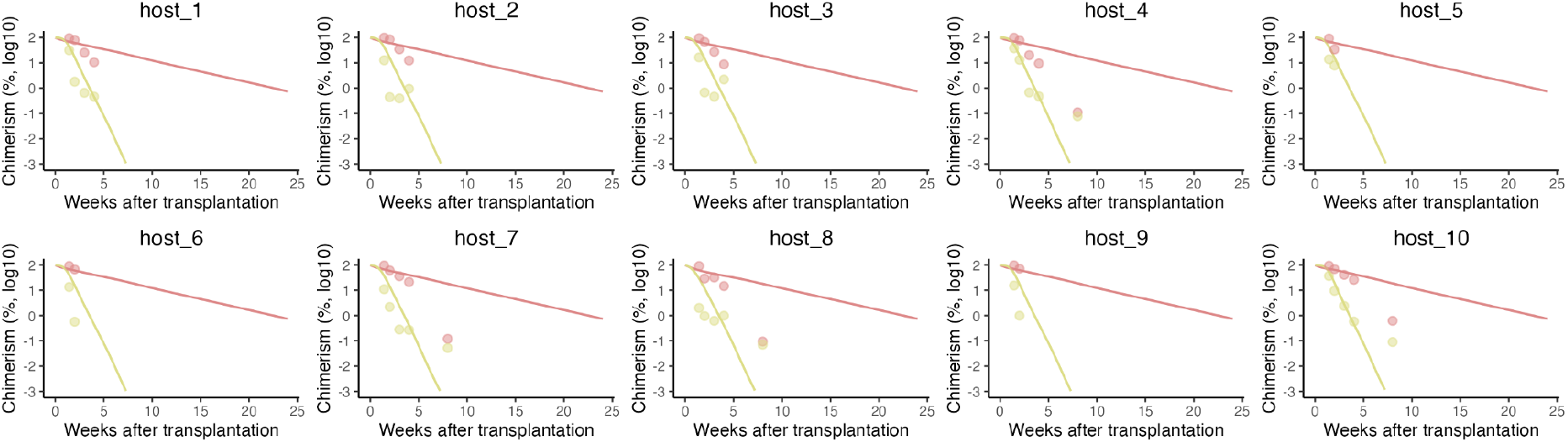
Individual recipient chimerism data in the bone marrow cell transplantation assay and best-fit model. Individual recipient chimerism data from the bone marrow cell transplantation assay (dots) and expected values of the mathematical model calculated with best-fit parameters (lines). Dots and lines in red and yellow correspond to erythrocytes and platelets, respectively.

**Figure S4.**
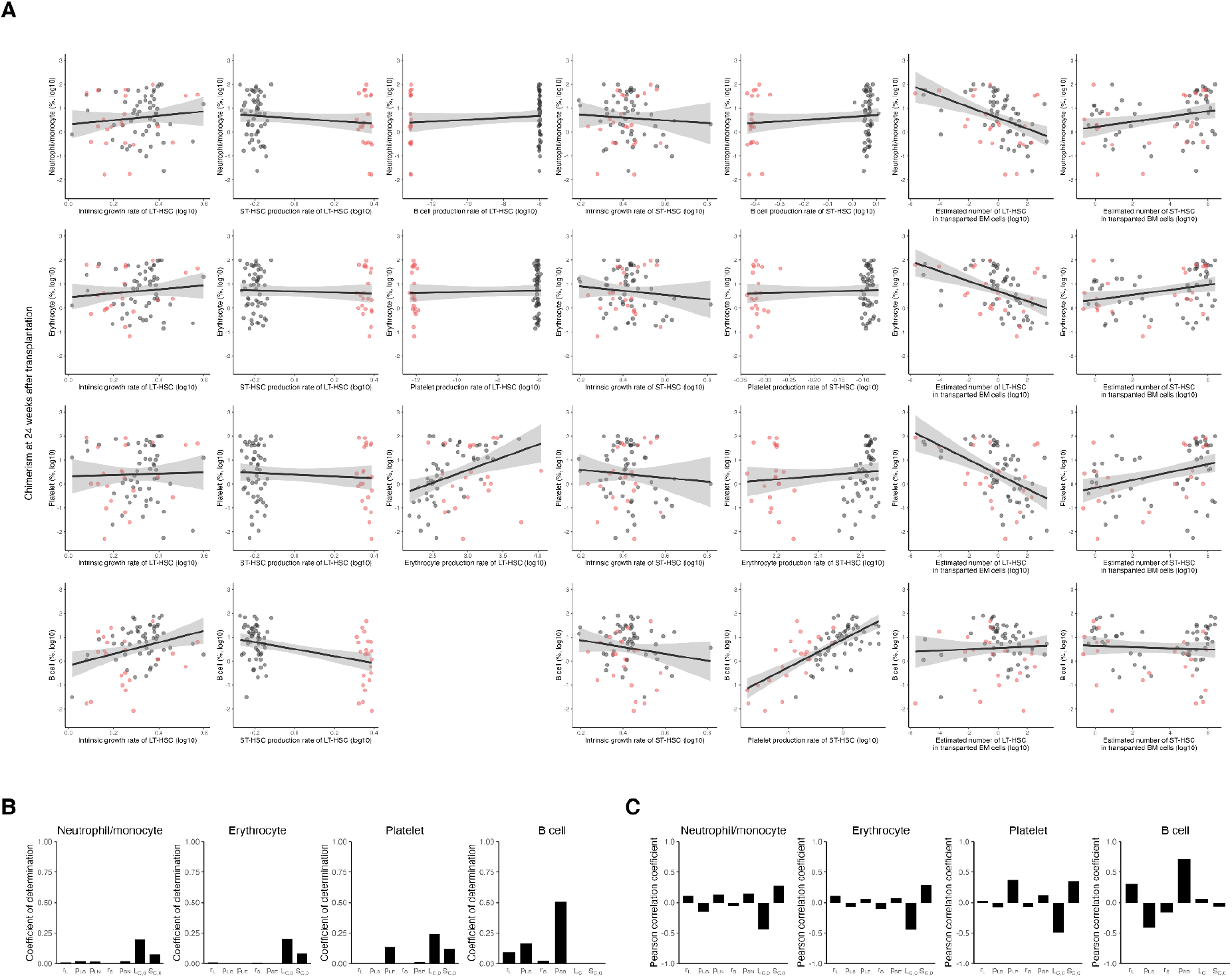
Comparison of estimated model parameters with experimentally obtained chimerism data. **(A)** Correlation between parameters in the mathematical model estimated from time-course data of chimerism in the single-cell transplantation assay and chimerism values of the relevant blood cell lines at 24 weeks after transplantation. Black and red dots correspond to the experiments in which young and aged HSCs were transplanted. Lines and shaded areas were obtained by linear regression. **(B-C)** Coefficient of determination **(B)** and Pearson correlation **(C)** coefficient in **(A)**.

**Figure S5.**
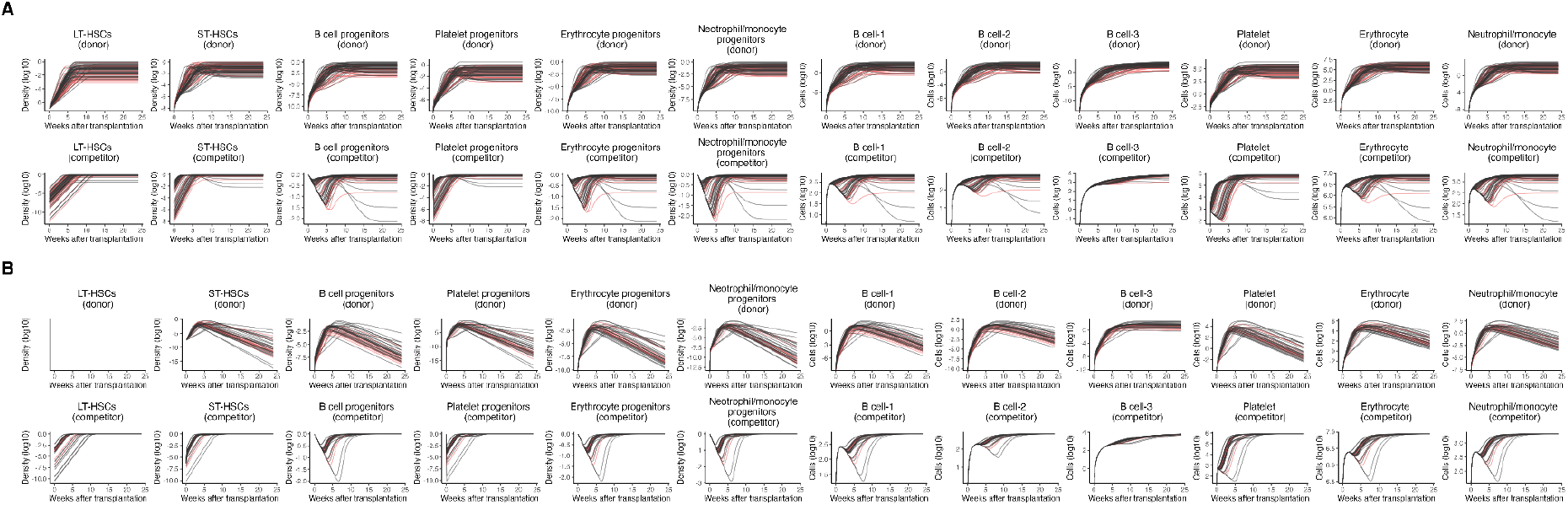
Expected density and number of each cell population in the single-cell transplantation assay. **(A-B)** Individual densities and numbers of each cell population in the single-cell transplantation of LT-HSC **(A)** and ST-HSC **(B)** calculated by the mathematical model with best-fit parameters. Black and red lines correspond to experiments in which young and aged HSCs were transplanted, respectively.

**Figure S6.**
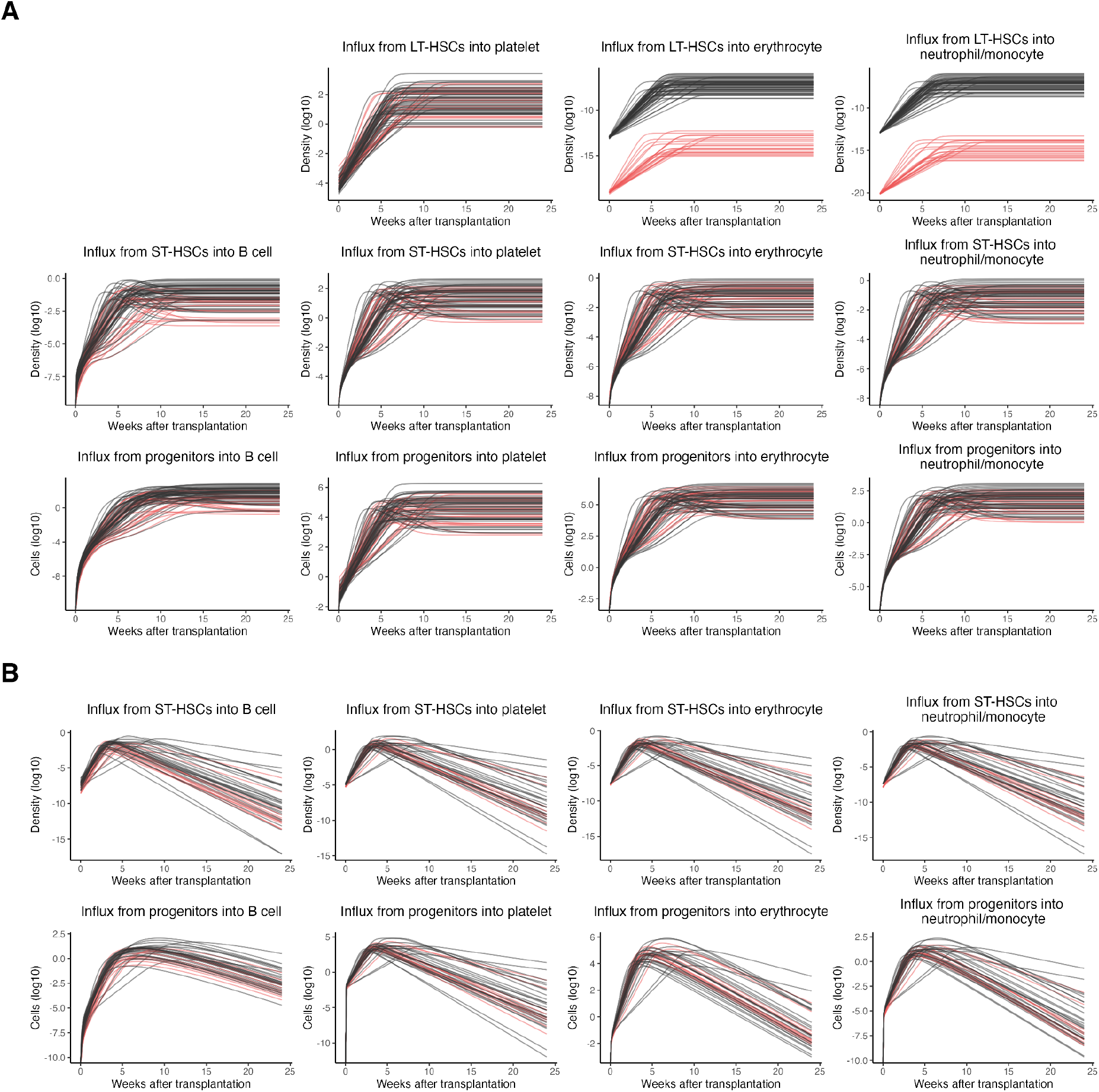
Expected influx of each cell population in the single-cell transplantation assay. **(A-B)** Production from the upper cell populations to each cell population at each time after transplantation calculated by the mathematical model with best-fit parameters (i.e., influx from LT-HSC into progenitors without going through ST-HSC, *p*_*Li,D,k*_ *L*_*i,D,k*_ (*t*) (*i* = *N, E, P*, and *k* = *n*_*L*1_, …, *n*_*L*75_, *n*_*S*1_, …, *n*_*S*39_), influx from ST-HSC to progenitors, *p*_*Si,D,k*_ *S*_*i,D,k*_ (*t*) (*i* = *N, E, P, B* and *k* = *n*_*L*1_, …, *n*_*L*75_, *n*_*S*1_, …, *n*_*S*39_), and influx from progenitors into mature cells, *α*_*i*_*d*_*i*_*Y*_*i,D,k*_ (*t*) (*i* = *N, E, P* and *k* = *n*_*L*1_, …, *n*_*L*75_, *n*_*S*1_, …, *n*_*S*39_) and *d*_*B*_*E*_*B*2,*D,k*_(*t*) (*k* = *n*_*L*1_, …, *n*_*L*75_, *n*_*S*1_, …, *n*_*S*39_)). Black and red lines correspond to the experiments in which young and aged HSCs were transplanted, respectively.

**Figure S7.**
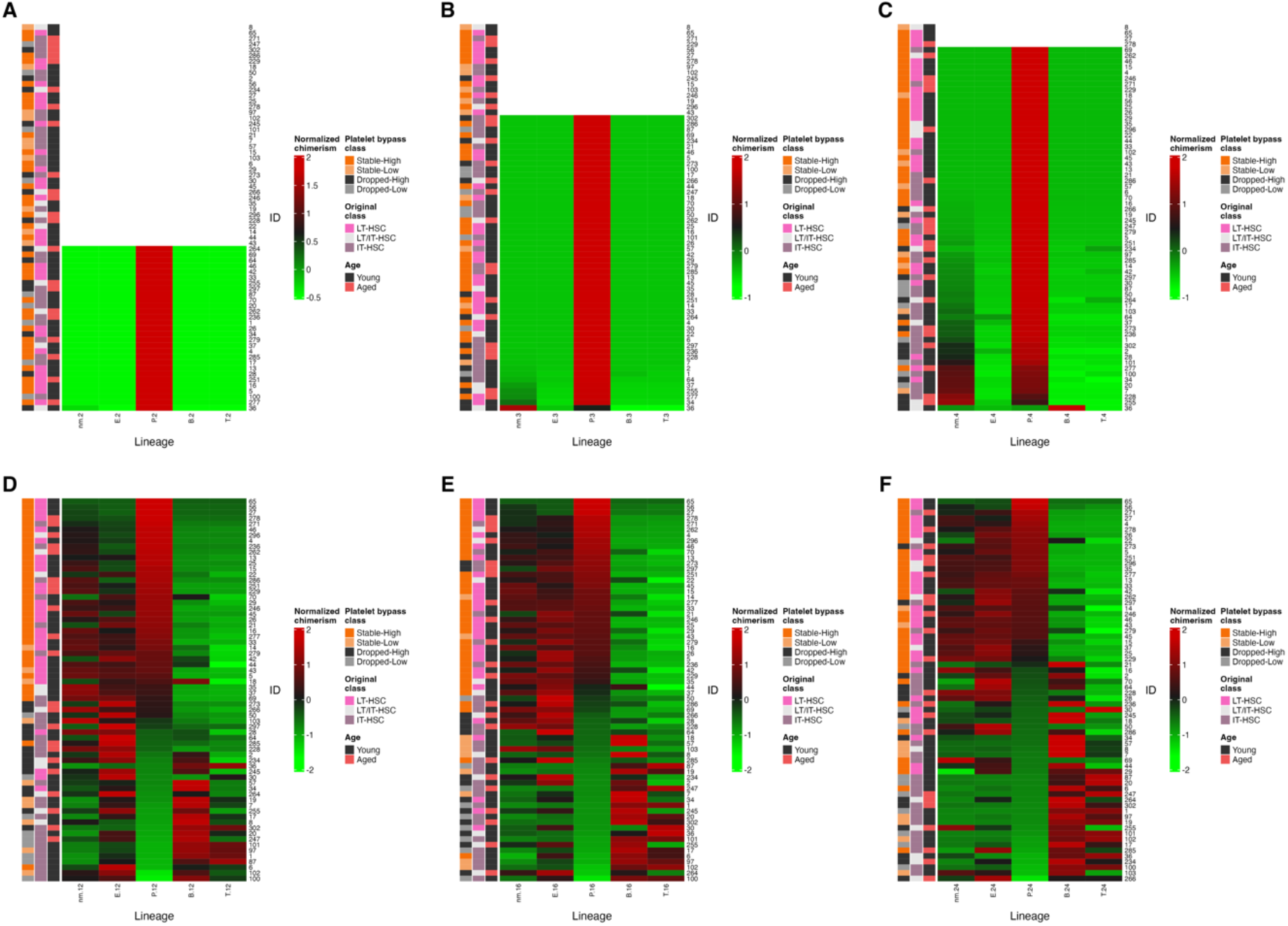
Normalized chimerism at 2-24 weeks after transplantation. **(A-F)** Normalized chimerism at 2 **(A)**, 3 **(B)**, 4 **(C)**, 12 **(D)**, 16 **(E)**, and 24 **(F)** weeks after transplantation with HSCs classified as LT-HSC or IT-HSC. The observations in which chimerism in all lineages were below the detection limit are indicated by white. The individuals are sorted in decreasing order using the normalized chimerism value of the platelet. The class defined by P-bypass, the class defined by the duration of reconstitution after transplantation, and age for transplanted HSCs are shown next to the heatmap.

**Figure S8.**
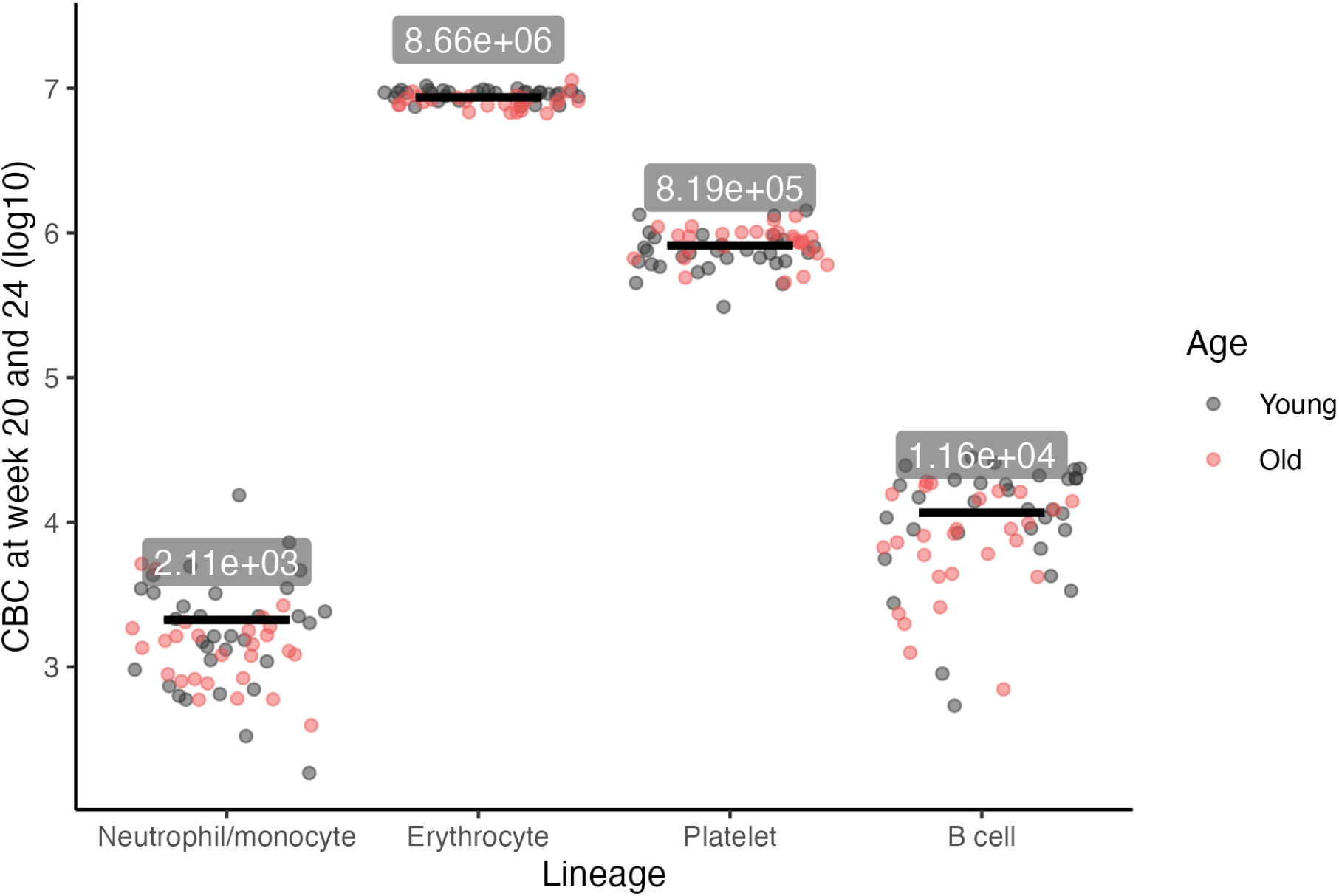
Measured CBC in single-cell transplantation assays. CBC values (dots) and their means (lines) for each blood cell lineage at 20 and 24 weeks after transplantation in single-cell transplantation assays in which CBCs were measured. The means were calculated by pooling the two time points and the types of young and aged HSCs.

### Supplementary Tables

**Table S1.** Estimated population parameters in nonlinear mixed-effect model.

| Name | Symbol | Unit | Fixed effect | Random effect<br>($\omega$) | Covariate<br>for age<br>($\beta_{age=aged}$) | Probability<br>distribution |
| --- | --- | --- | --- | --- | --- | --- |
| Maximum<br>amount of<br>LT-HSCs<br>(Carrying<br>capacity of<br>LT-HSC) | $L_s$ | – | $8.42 \times 10^6$ | – | – | – |
| | $L^* (= L_s + 1)$ | – | $8.42 \times 10^6$<br>(calculated) | – | – | – |
| Maximum<br>amount of<br>ST-HSCs | $S_s$ | – | $2.98 \times 10^7$ | – | – | – |
| | $S^* (= S_s + 1)$ | – | $2.98 \times 10^7$<br>(calculated) | – | – | – |
| Carrying<br>capacity of<br>ST-HSCs | $K_s \left( = \frac{r_{s,c}}{p_{LS,c} + r_{s,c}} \right)$ | – | $2.98 \times 10^7$<br>(calculated) | – | – | – |
| Intrinsic<br>growth rate<br>of LT-HSCs | $r_{L,c}$ | week<sup>-1</sup> | 2.41 | – | – | – |
| | $r_{L,d}$ | – | 1<br>(fixed) | 0.305 | –0.303 | log-normal |
| | $r_{L,D} (= r_{L,c}r_{L,d})$ | week<sup>-1</sup> | 2.41<br>(calculated) | – | – | – |
| Intrinsic<br>growth rate<br>of ST-HSCs | $r_{s,c}$ | week<sup>-1</sup> | 2.75 | – | – | – |
| | $r_{s,d}$ | – | 1<br>(fixed) | 0.479 | 0.105 | log-normal |
| | $r_{s,D} (= r_{s,c}r_{s,d})$ | week<sup>-1</sup> | 2.75<br>(calculated) | – | – | – |
| ST-HSC<br>production<br>rate from<br>LT-HSCs | $p_{LS,c}$ | week<sup>-1</sup> | 0.649 | – | – | – |
| | $p_{LS,d}$ | – | 1<br>(fixed) | 0.314 | 1.27 | log-normal |
| | $p_{LS,D} (= p_{LS,c}p_{LS,d})$ | week<sup>-1</sup> | 0.649<br>(calculated) | – | – | – |
| Neutrophil/<br>monocyte<br>production<br>rate from | $p_{LN,c}$ | week<sup>-1</sup> | $1.04 \times 10^{-6}$ | – | – | – |
| | $p_{LN,d}$ | – | 1<br>(fixed) | 2.90 | –16.6 | log-normal |
| LT-HSCs | $p_{LN,D} (= p_{LN,C}p_{LN,d})$ | week<sup>-1</sup> | $1.04 \times 10^{-6}$<br>(calculated) | – | – | – |
| Erythrocyte<br>production<br>rate from<br>LT-HSCs | $p_{LE,C}$ | week<sup>-1</sup> | $9.41 \times 10^{-7}$ | – | – | – |
| | $p_{LE,d}$ | – | 1<br>(fixed) | 5.30 | –14.0 | log-normal |
| | $p_{LE,D} (= p_{LE,C}p_{LE,d})$ | week<sup>-1</sup> | $9.41 \times 10^{-7}$<br>(calculated) | – | – | – |
| Platelet<br>production<br>rate from<br>LT-HSCs | $p_{LP,C}$ | week<sup>-1</sup> | 413 | – | – | – |
| | $p_{LP,d}$ | – | 1<br>(fixed) | 1.31 | 1.01 | log-normal |
| | $p_{LP,D} (= p_{LP,C}p_{LP,d})$ | week<sup>-1</sup> | 413<br>(calculated) | – | – | – |
| Neutrophil/<br>monocyte<br>production<br>rate from<br>ST-HSCs | $p_{SN,C}$ | week<sup>-1</sup> | 1.19 | – | – | – |
| | $p_{SN,d}$ | – | 1<br>(fixed) | 0.222 | –1.13 | log-normal |
| | $p_{SN,D} (= p_{SN,C}p_{SN,d})$ | week<sup>-1</sup> | 1.19<br>(calculated) | – | – | – |
| Erythrocyte<br>production<br>rate from<br>ST-HSCs | $p_{SE,C}$ | week<sup>-1</sup> | 0.813 | – | – | – |
| | $p_{SE,d}$ | – | 1<br>(fixed) | 0.166 | –0.515 | log-normal |
| | $p_{SE,D} (= p_{SE,C}p_{SE,d})$ | week<sup>-1</sup> | 0.813<br>(calculated) | – | – | – |
| Platelet<br>production<br>rate from<br>ST-HSCs | $p_{SP,C}$ | week<sup>-1</sup> | 417 | – | – | – |
| | $p_{SP,d}$ | – | 1<br>(fixed) | 0.492 | –0.969 | log-normal |
| | $p_{SP,D} (= p_{SP,C}p_{SP,d})$ | week<sup>-1</sup> | 417<br>(calculated) | – | – | – |
| B cell<br>production<br>rate from<br>ST-HSCs | $p_{SB,C}$ | week<sup>-1</sup> | 0.730 | – | – | – |
| | $p_{SB,d}$ | – | 1<br>(fixed) | 1.19 | –1.61 | log-normal |
| | $p_{SB,D} (= p_{SB,C}p_{SB,d})$ | week<sup>-1</sup> | 0.730<br>(calculated) | – | – | – |
| Death rate of<br>neutrophils/ | $\delta_{N,D}, \delta_{N,C}$ | week<sup>-1</sup> | 0.512 | – | – | – |
| monocytes |  |  |  |  |  |  |
| Death rate of erythrocytes | $\delta_{E,D}, \delta_{E,C}$ | week<sup>-1</sup> | 0.493 | – | – | – |
| | $\delta_{E,R}$ | week<sup>-1</sup> | 0.204 | – | – | – |
| Death rate of platelets | $\delta_{P,D}, \delta_{P,C}$ | week<sup>-1</sup> | 0.624 | – | – | – |
| | $\delta_{P,R}$ | week<sup>-1</sup> | 1.62 | – | – | – |
| Death rate of B cells | $\delta_{B,D}, \delta_{B,C}$ | week<sup>-1</sup> | 0.0433 | – | – | – |
| Differentiation rate of neutrophil/monocyte progenitors | $d_N (= p_{LN,C} + p_{SN,C})$ | week<sup>-1</sup> | 1.19<br>(calculated) | – | – | – |
| Differentiation rate of erythrocyte progenitors | $d_E (= p_{LE,C} + p_{SE,C})$ | week<sup>-1</sup> | 0.813<br>(calculated) | – | – | – |
| Differentiation rate of platelet progenitors | $d_P (= p_{LP,C} + p_{SP,C})$ | week<sup>-1</sup> | 830<br>(calculated) | – | – | – |
| Differentiation rate of B cell progenitors | $d_B (= p_{SB,C})$ | week<sup>-1</sup> | 0.730<br>(calculated) | – | – | – |
| The number of neutrophils/monocytes in steady-state | $Z_N^*$ | cells | $2.11 \times 10^3$<br>(fixed) | – | – | – |
| The number of erythrocytes in steady-state | $Z_E^*$ | cells | $8.66 \times 10^6$<br>(fixed) | – | – | – |
| The number of platelets in steady-state | $Z_P^*$ | cells | $8.19 \times 10^5$<br>(fixed) | – | – | – |
| The number of B cells in | $Z_B^*$ | cells | $1.16 \times 10^4$<br>(fixed) | – | – | – |
| steady-state |  |  |  |  |  |  |
| Expansion rate of neutrophils/monocytes | $\alpha_N \left( = \frac{\delta_{N,C} Z_N^*}{d_N} \right)$ | cells | $1.04 \times 10^9$<br>(calculated) | — | — | — |
| Expansion rate of erythrocytes | $\alpha_E \left( = \frac{\delta_{E,C} Z_E^*}{d_E} \right)$ | cells | $4.54 \times 10^{12}$<br>(calculated) | — | — | — |
| Expansion rate of platelets | $\alpha_P \left( = \frac{\delta_{P,C} Z_P^*}{d_P} \right)$ | cells | $1.66 \times 10^3$<br>(calculated) | — | — | — |
| Expansion rate of B cells | $\alpha_B \left( = \frac{\delta_{B,C} Z_B^*}{d_B} \right)$ | cells | 688<br>(calculated) | — | — | — |
| The amount of LT-HSCs in competitor cells | $L_{C,0}$ | — | 5.66 | 5.72 | — | log-normal |
| The amount of ST-HSCs in competitor cells | $S_{C,0}$ | — | 625 | 7.68 | — | log-normal |
| Variance of residual errors | $a_{F,N}$ | — | 1.99 | — | — | — |
| | $a_{F,E}$ | — | 1.52 | — | — | — |
| | $a_{F,P}$ | — | 2.27 | — | — | — |
| | $a_{F,B}$ | — | 1.61 | — | — | — |
| | $a_{T,N}$ | — | 0.743 | — | — | — |
| | $a_{T,E}$ | — | 0.179 | — | — | — |
| | $a_{T,P}$ | — | 0.592 | — | — | — |
| | $a_{T,B}$ | — | 1.76 | — | — | — |

**Table S2. Estimated individual parameters in nonlinear mixed-effect model**

*id, class, age, individual model parameter values are listed in TableS2.csv.*

### Supplementary Text

#### HSC differentiation model

To analyze the dynamics of chimerism in the single-cell transplantation assays, we developed a differentiation model of HSCs. In the base model, self-renewing LT-HSCs differentiate into ST-HSCs, which then differentiate into mature cells through progenitors. Using ordinary differential equations, the mathematical model is described as follows:

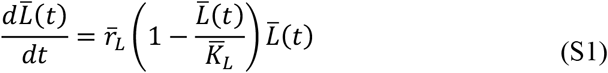

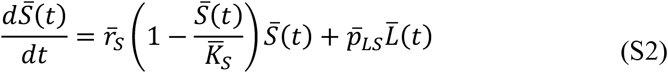

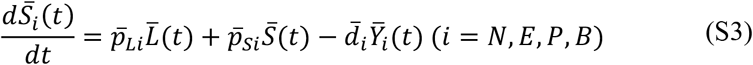

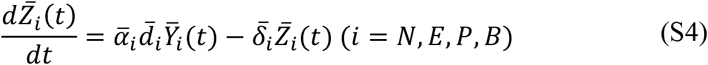

The variables 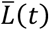 and 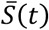 represent the number of LT-HSCs and ST-HSCs at time *t*, and the numbers of progenitors and mature cells at time *t* in neutrophils/monocytes (*i* = *N*), erythrocytes (*i* = *E*), platelets (*i* = *P*), and B cells (*i* = *B*) are represented by 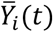 and 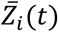, respectively. The ability of LT-HSCs and ST-HSCs to proliferate by self-replication is described as logistic proliferation using the intrinsic growth rate, 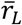 and 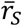, and the carrying capacity 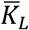 and 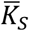, respectively. ST-HSCs are produced from LT-HSCs at rate 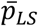, and progenitor cells are produced from ST-HSCs and ST-HSCs at rates 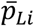 and 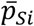 (*i* = *N, E, P, B*), respectively. Direct production of progenitor cells from LT-HSCs, 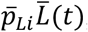, represents a bypass pathway with lineage-restricted differentiation. Here 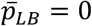 because we consider only the bypass pathway for differentiation into myeloid lineages. Progenitors differentiate at rate 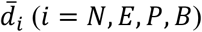 and mature cells are produced with expansion rate 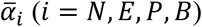. Mature cells die at rate 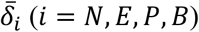.

In normal blood cell production, all populations are considered to be in steady-state, which is described as 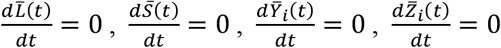 and 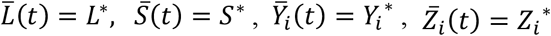, where 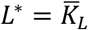. Using these relationships, **Eqs. (S1-S4)** can be transformed as follows:

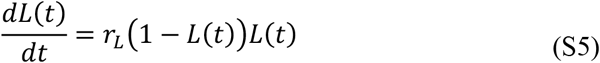

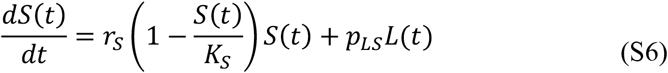

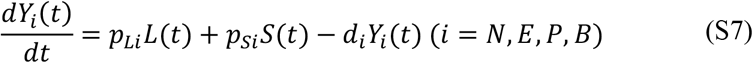

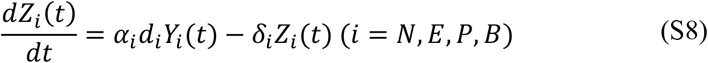

Here, the variables are converted as 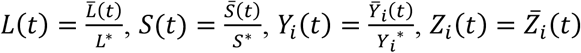. Thus, the parameters are converted as 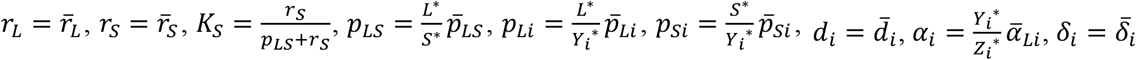. The steady-state relationship allows us to treat some parameters as *d*_*i*_ = *p*_*Li*_ + *p*_*Si*_ and 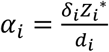.

### Mathematical model for single-cell transplantation

In the single-cell transplantation assay, BM cells (competitors) are transplanted at the same time as single cells (donor cells). In addition, recipient cells are included in the measurement of erythrocytes and platelets. The competitive proliferation of donor- (*j* = *D*) and competitor- (*j* = *C*) and recipient- (*j* = *R*) derived cells in the single-cell transplantation assay can be expressed by extending **Eqs. (S5-S8)** as follows:

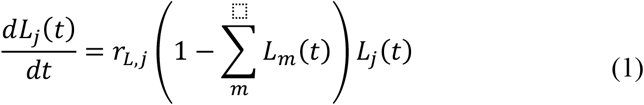

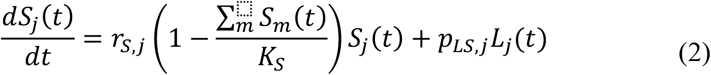

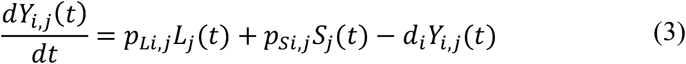

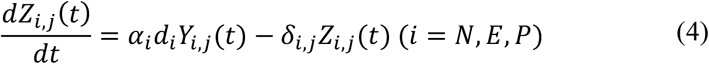

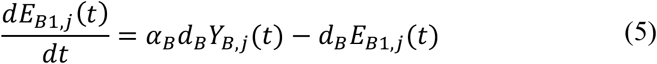

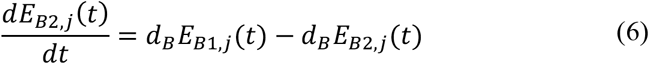

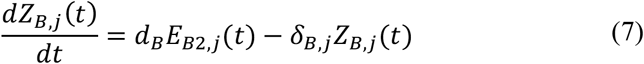

where *t* is the time after transplantation. The subscript *j* in each variable and parameter corresponds to the cell origin (donor cells, competitors, recipient cells). Here, we assume the existence of an eclipse phase in the differentiation of progenitor cells to mature cells to represent the slow production of B cells. This delay is explained by the commonly used linear chain trick. Since the recipient mice have been lethally irradiated and are not capable of producing blood cells, the parameters other than the death rate of mature cells, *δ*_*i,j*_, related to the recipient-derived cells are zero. The differentiation of non-HSC populations, which are described by parameters *d*_*i*_ and *α*_*i*_, is assumed to be the same for donor- and recipient-derived cells. Death of mature cells was assumed to be different only for recipient-derived cells as an effect of irradiation, *δ*_*i,D*_ = *δ*_*i,C*_.

Chimerism and CBC were measured in the single-cell transplantation assay. In the measurements, donor cells are identified by KuO, a propensity protein. In neutrophils/monocytes and B cells, competitors and recipient cells are distinguished by the expression of Ly5.1 and Ly5.2, which are cell surface markers of lymphocytes, but not in erythrocytes and platelets. Thus, the chimerism, *F*_*j*_(*t*), and CBC, *T*_*j*_(*t*), for each lineage obtained in the experiment are calculated using **Eqs. (1-7)** as follows:

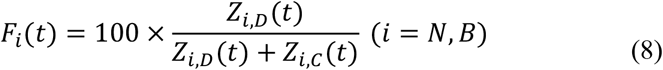

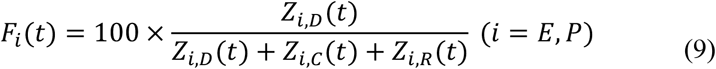

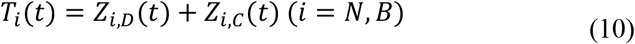

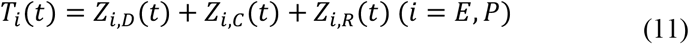

In the single-cell transplantation assay, single cells in pHSCs are transplanted. The initial conditions of donor cells are *L*_*D*_(0) = 1/*L*^*^, *S*_*D*_(0) = 0, *Y*_*i,D*_ (0) = 0, *E*_*B*1,*D*_ (0) = 0, *E*_*B*2,*D*_(0) = 0, *Z*_*i,D*_(0) = 0 or *L*_*D*_(0) = 0, *S*_*D*_ (0) = 1/*S*^*^, *Y*_*i,D*_(0) = 0,*E*_*B*1,*D*_ (0) = 0, *E*_*B*2,*D*_ (0) = 0, *Z*_*i,D*_ (0) = 0 for the transplantations in which transplanted single cells are classified as LT-HSCs (including IT-HSCs) or ST-HSCs, respectively. Assuming that the 2 × 10^5^ BM cells transplanted as competitor cells contain a small amount of HSCs and progenitor cells that can be responsible for blood cell production, the initial condition was set to *L*_*C*_(0) = *L*_*C*,0_, *S*_*C*_ (0) = *S*_*C*,0_, *Y*_*i, C*_ (0) = 1,*E*_*B*1,*C*_ (0) = 0, *E*_*B*2,*C*_ (0) = 0, *Z*_*i,C*_ (0) = 0. Since recipient mice were lethally irradiated, we adopted initial conditions, *L*_*R*_(0) = 0, *S*_*R*_(0) = 0, *Y*_*i,R*_ (0) = 0,*E*_*B*1, *R*_ (0) = 0, *E*_*B*2, *R*_ (0) = 0, *Z*_*i,R*_ (0) = *Z*_*i*_^*^, in which HSCs and progenitor cells were absent and assumed that they were unable to produce blood cells. The numbers of mature cells derived from recipient cells, *Z*_*i*_ ^*^, were estimated as averages of the values of CBC at 20 and 24 weeks after transplantation (**Figure S8**).

### Mathematical model for BM cell transplantation

To quantify the kinetics of recipient-derived cells of erythrocytes and platelets, an experiment was performed in which BM cells labeled with KuO were transplanted. In this experiment, there are cells derived from BM cells, corresponding to the competitors of the single-cell transplantation assay, and recipient-derived cells. Chimerism was measured as the percentage of recipient-derived erythrocytes and platelets in PB, which was described using **Eqs. (1-7)** as follows:

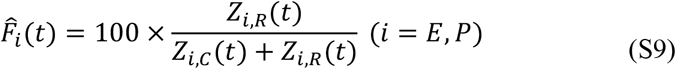

Assuming that the composition of the BM cells to be transplanted is similar to that of the competitors in the single-cell transplantation assay, the initial values for **Eqs. (1-7)** are *L*_*D*_(0) = 0, *S*_*D*_ (0) = 0, *Y*_*i,D*_ (0) = 0, *E*_*B*1,*D*_ (0) = 0, *E*_*B*2,*D*_ (0) = 0, *Z*_*i,D*_ (0) = 0, *L*_*C*_ (0) = *L*_*C*,0_, *S*_*C*_ (0) = *S*_*C*,0_, *Y*_*i,C*_ (0) = 1, *E*_*B*1,*C*_ (0) = 0, *E*_*B*2,*C*_ (0) = 0, *Z*_*i,C*_ (0) = 0, *L*_*R*_ (0) = 0, *S*_*R*_ (0) = 0, *Y*_*i,R*_ (0) = 0, *E*_*B*1,*R*_ (0) = 0, *E*_*B*2,*R*_ (0) = 0, *Z*_*i,R*_ (0) = *Z*_*i,R*_ (0) = *Z*_*i*_^*^ in BM cell transplantation.

### Model parameter estimation

A nonlinear mixed-effect model was used to fit the mathematical model for chimerism and CBC given by **Eqs. (8-11)** and **(S9)** to the dataset from the single-cell and BM cell transplantation assay. Assume that all parameters of differentiation kinetics of competitors are normal values of blood cell production (i.e., *r*_*L,C*_ = *r*_*L*_, *r*_*S,C*_ = *r*_*S*_, *p*_*LS,C*_ = *p*_*LS*_, *p*_*Li,C*_ = *p*_*Li*_, *p*_*Si,C*_ = *p*_*Si*_). In addition, assuming that the transplanted single cells are heterogeneous in their proliferation and replication abilities, we further assumed that the values of their parameters vary around normal values. By introducing new scaling parameters with subscript d, the parameter for the differentiation kinetics of HSCs in donor cells is expressed as *r*_*L,D*_ = *r*_*L,C*_ *r*_*L,d*_, *r*_*S,D*_ = *r*_*S,C*_ *r*_*S,d*_, *p*_*LS,D*_ = *p*_*LS,C*_ *p*_*LS,d*_, *p*_*Li,D*_ = *p*_*Li,C*_ *p*_*Li,d*_, *p*_*Si,D*_ = *p*_*Si,C*_ *p*_*Si,d*_. If the scaling parameters for individual *k* follow log-normal distributions and have differences between young and aged HSCs, then the parameters representing the ability of LT-HSCs, *ψ*_*L,j*_ = {*r*_*L,j*_, *p*_*LS,j*_, *p*_*Li,j*_}(*j* = *D, C*) and ST-HSCs, *ψ*_*S,j*_ = {*r*_*S,j*_, *p*_*Si, j*_} (*j* = *D, C*), for individual *k* follow the following formula:

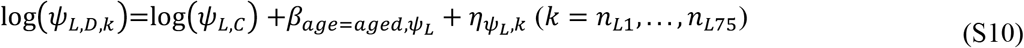

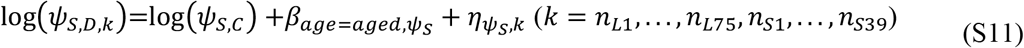

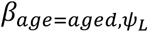 and 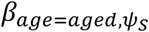 are covariates for the ability of aged HSCs. 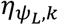 and 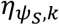 are random effects following normal distributions with mean 0 and standard deviations 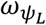 and 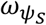. The individual IDs of single cells classified as LT-HSC and ST-HSC and mice to which the single cells were transplanted in the single-cell transplantation are denoted as *n*_*L*,1_, …, *n*_*L*,75_ and *n*_*S*,1_, …, *n*_*S*,39_, respectively. Since the number of LT-HSCs and ST-HSCs in competitors is unknown and the number in transplanted BM cells is expected to be heterogeneous, their distribution follows the following formula:

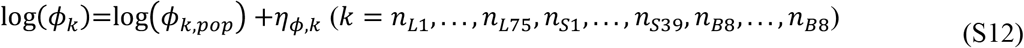

*η*_*ϕ,k*_ is a random effect following normal distributions with mean 0 and standard deviations *ω*_*ϕ,k*_. The individual IDs of mice in the BM transplantation are denoted as *n*_*B*1_, …, *n*_*B*8_.

The parameters of the model were estimated by fitting the mathematical model to the corresponding data, assuming the following observation model:

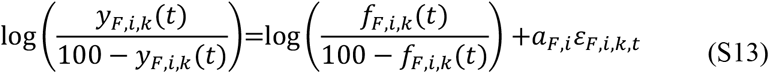

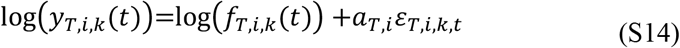

where,

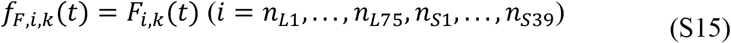

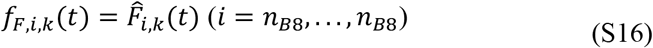

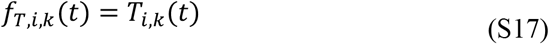

*ε*_*F,i,k,t*_ and *ε*_*T,i,k,t*_ are observation errors for chimerism and CBC data, *y*_*F,i,k*_ (*t*) and *y*_*T,i,k*_ (*t*), of lineage *i* in individual *k* observed at time *t*, which follows a standard normal distribution. *a*_*F,i*_ and *a*_*T,i*_ define variances of observation errors for chimerism and CBC. To account for the data points under the detection limit (detection limit is 0.001% for chimerism), the likelihood function assumed the data under the detection limit are censored. The estimated population parameters and individual parameters are listed in **Table S1** and **S2**, respectively.

